# No pervasive size trend in global community dynamics

**DOI:** 10.1101/2021.05.11.443574

**Authors:** J. Christopher D. Terry, Jacob D. O’Sullivan, Axel G. Rossberg

## Abstract

While recent meta-analyses have suggested that local taxonomic richness has on average remained invariant, potential shifts in functional traits remain underexplored at global scales. Here, by linking the largest cross-taxa community time series database to multiple trait databases, we show that within communities there is no clear trend between size traits and changes in abundance rank over time. This suggests that there is no pervasive tendency across biomes for larger species to be doing proportionally better or worse than smaller species.

---

Recent analyses have found that despite high and increasing levels of community turnover, there is no clear overall trend in the local-scale species richness^1–4^. However, it remains unclear how this result translates into functional changes. One of the most fundamental functional traits of a species is its size^5,6^ and there is an expectation that a warming climate is will likely favour smaller bodysizes^7,8^. Further, larger species have been more extinction prone during some previous mass-extinctions^9,10^, are more likely to show strong recent population declines^11^, and (although relationships are threat-dependent) tend to be assessed at a higher risk of extinction^12^ due to longer generational intervals and increased threat from habitat fragmentation and hunting^13^.

One might therefore expect a detectable signal of shifts in community trait values beneath the apparent underlying consistency in taxonomic diversity. To examine this, we tested whether the body size of a species is correlated to a relative change in abundance through time using the publicly available BioTIME database^14^. This database is the largest collection of time series of ecological communities and has wide geographic and taxonomic scope^14^. After cleaning and standardising the names associated with the records, we linked six fundamental ‘size’ traits from four openly accessible trait databases representing four guilds – adult body mass from a database of amniote life history traits^15^, adult body length and qualitative body size of marine species from the WoRMS database^16^, plant maximum height and seed mass from the TRY database^17^, and maximum body length of fish from a compilation^18^ based on data in the FishBase repository^19^.

Observations from single-location studies were collated, while widely dispersed studies were separately binned into a global grid of cells, each approximately 10km wide, and data from each study and cell were treated as discrete assemblages, following previous analyses^1,20^. Selecting only assemblages with quantified observations of ≥10 species, over ≥5 years, and with ≥40% of the species having records for at least one trait, we generated 22,915 community time series from 167 studies. This filtered dataset represented 2,516,175 observations of 12,033 species, of which 8,238 could be linked to at least one ‘size’ trait (representing 84.6% of observations). Equally weighting studies, the average time series length was 17.2 years (range 5-70.5) and the average number of species per included assemblage was 56.7 (range 10-337).

To focus on changes within the community and mitigate the risk of issues with changing sampling intensities through time, for each trait and community assemblage time series for which there was sufficient data, we calculated ‘τ’ the Kendall rank correlation coefficient between the trait in question and change in relative-rank of the abundance of each species within the assemblage (Fig 1a). This gives a non-parametric measure of whether larger species are more or less likely than smaller species to have increased their relative rank through time and, importantly, can be calculated where trait values for only a fraction of the observed species are available. To weight each study within BioTIME equally, where there were multiple assemblages per study these were averaged to generate a τ value for each possible study-trait combination. In order to provide a null-model against which to evaluate the statistical significance of this multistage analysis, we repeated the procedure with 100 trait randomisations per assemblage.

**Figure 1.**
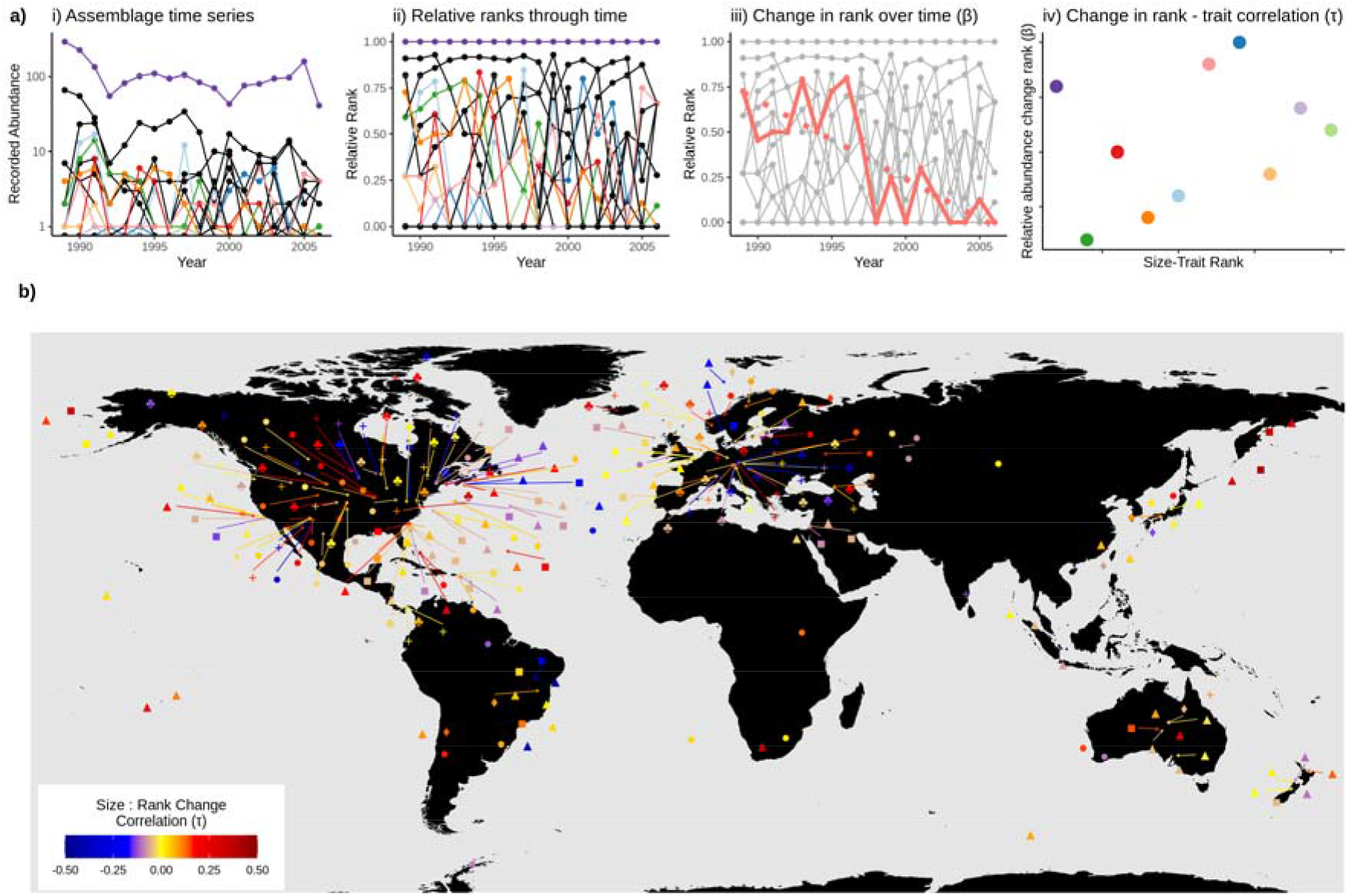
a) Example of the non-parametric approach taken to accomadate patchy trait data and possible variation in sampling intensity through time for an arbitrary example assemblage. Here species with trait values are coloured, others without trait data are black. (i) Quantitative survey data is combined to generate assemblage time series. (ii) the relative ranks of each species are calculated for each year, through which is fit a linear model summarising the rate of change (β) in a species rank through time (iii). For those species where trait data is available, the concordance between body size traits and abundance rank changes (iv) is then summarised by τ, the Kendal rank-correlation coefficient (here, τ = 0.24). b) Global distribution of studies, for each tested trait, showing average τ for each study-trait combination. Marine: Body length = ▲, Marine: Qualitative body size = ■, Fish: Maximum length = ⍰, Amniotes: Adult body mass = ⍰, Plants: Seed mass = +, Plants: Maximum height = ⍰. Note that the locations are shown as centre point of each study, which can cause oceanic studies to be ‘located’ on land. See Extended Data Table 4 for full details of study-level results.

For five of the six tested size traits the distribution of study τ values was not significantly different from the null model (Fig. 2, Extended Data Table 1). The one exceptional trait (amniote body mass) showed an overall average positive relationship between size and relative abundance trends. Possible confounding factors for the value of τ associated with each study, namely the total span of the time series, the number of sample points, the species richness, the average trait completeness, the number of assemblages within the study, the grain of the study and the absolute latitude did not consistently predict τ (Extended Data Figure 1, Tables 2-3).

**Figure 2.**
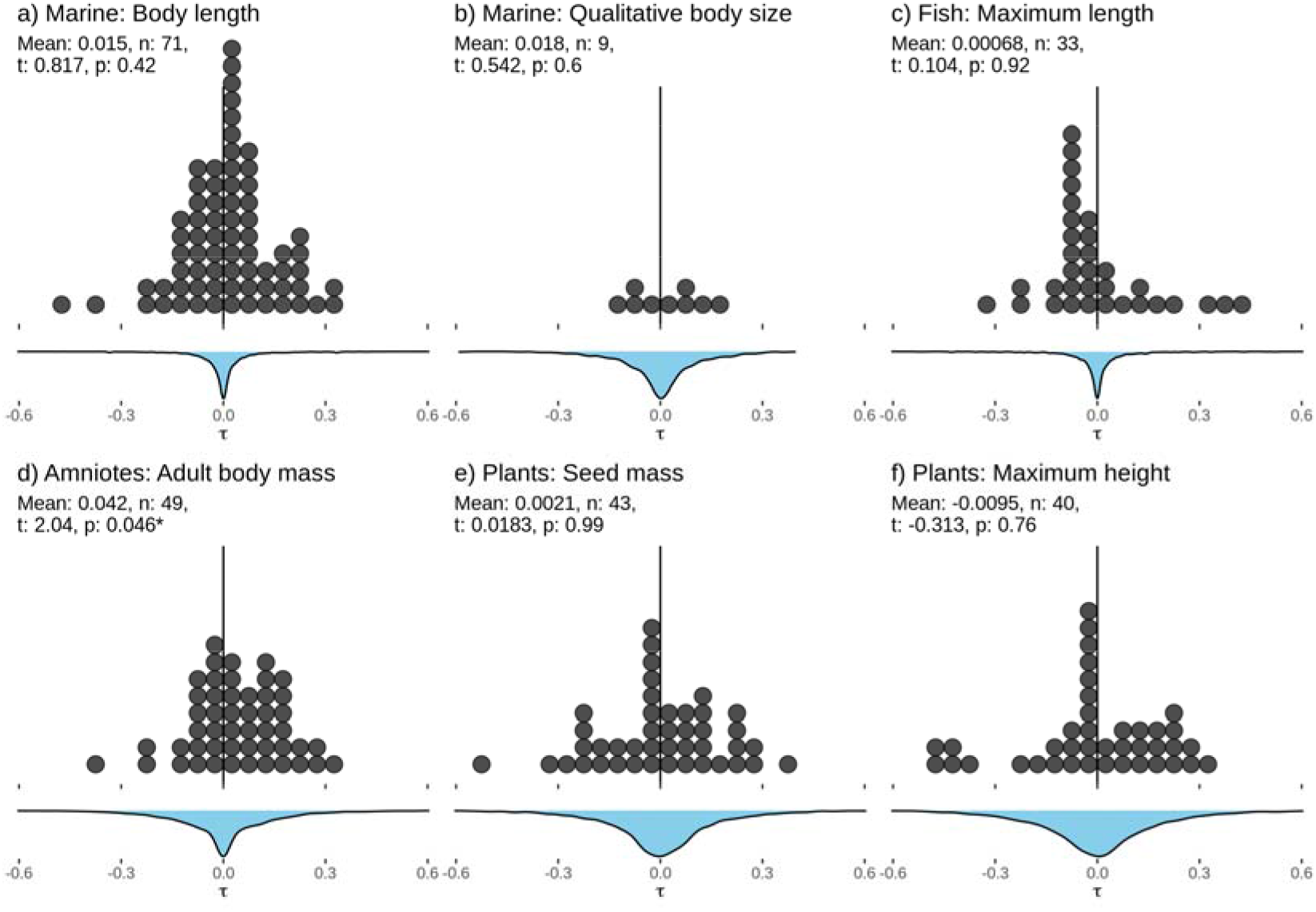
Distribution of Kendall’s rank correlation coefficient between the 6 tested body-size traits and changes in relative abundance through time (‘τ’). Each dot represents one study, averaging across the constituent assembly time series for studies of large spatial extent. Results are binned into classes 0.05 units of τ wide. Blue density plots show the distribution of 50 randomisations of available trait data. Statistical results refer to two-sided two-sample t-tests, comparing the distribution of observed τ values to the null distribution. Means shown are for the observed results. Full statistical results are given in Extended Data Table 1.

These results indicate that there is not yet evidence for widely pervasive trends in a core functional trait, body size. Importantly however, this study should not be seen as a refutation of the enhanced threats faced by the very largest apex species^21^. Rather, against a background of considerable turnover^2,3^ across the whole community assemblage, on average, species positions in communities are being taken up by species of comparable size. This suggests that previously identified shifts towards smaller species found in some systems^7,22^ may not be as universal as currently expected^8^ and aligns with the divergent changes in global body-size abundance distributions observed between mammal guilds^23^. However, we note that the variance in *τ* across the dataset was consistently larger than that generated by a null model randomising the available trait data within assemblages (Fig 2). This implies that many individual communities are experiencing significant directional change in their body size traits, rather than local stasis or indeed active regulation.

The overall positive association across the amniote studies could have a number of causes that would benefit from further investigation. One putative explanation is that anthropogenic dispersal limitations (generally considered to act more strongly against smaller species) may be having a greater immediate impact than climate change^24^. There are also indications of differences between terrestrial and marine systems. Previous work with the same datasets^1,20^ has found greater species richness and abundance changes in marine systems than in terrestrial systems, while here we see a signal of greater trait-changes in the (largely terrestrial) amniotes. In our study, the fish lengths trait displayed a distinctive distribution of τ (Fig 2c), with a modal peak of studies showing small negative values, and then a tail of strongly positive relationships. This guild is also the most likely to have experienced sustained anthropogenic pressure^25^ and many of the ‘fish’ datasets in BioTIME include data from surveys of actively fished and managed areas. Accurately quantifying marine community trends is a challenge^26,27^, but this distinctive pattern could be tentatively interpreted as a signal of the imposition or relaxation of anthropogenic pressure across marine systems^28,29^.

Our analysis necessarily sacrifices fine resolution for global scale. Although the lack of consistent study-level drivers of τ suggests that the results are unlikely to be solely determined by biases within the BioTIME database, future work should seek to improve the scope and resolution of available data to enable more strongly parametric analyses and examine additional measures of community change. While available trait databases of amniotes and fish are carefully curated, checked and taxonomically tidy, the other databases pose more problems in terms of taxonomic matching and consistency of trait measurements. Without direct correspondence between dynamics and trait data sources, it is necessary to take traits as fixed values for each species, despite known differences in traits in time^30–32^ and space^33^ that may themselves represent responses to global change. However, in Celtic sea fish within-species shifts have been shown to contribute less to community-level size shifts than changes in species composition ^34^. We also note that ‘size’ traits for indeterminately growing plants have a less clear meaning than for animals. However, both seed size and maximum height are linked to environmental variables^35,36^ and could therefore be hypothesised to influence species responses to global change.

Many of the criticisms and defences regarding earlier studies using the BioTIME dataset, and indeed other metanalyses of large collections of time series, also apply to this work^37^. The statistical approach was developed to be as robust as possible to both incomplete trait data and changes in overall abundance and sampling effort through time. However, it must be noted that our largely non-parametric approach could lack the power and resolution to identify subtle changes. Biases in the underlying BioTIME database towards vertebrate taxa and North American and European sites^14^ are further exaggerated when crossed with trait data availability (Figure 1b). One particularly concerning gap is the absence of any insect studies due to a paucity of usable trait information. Observations suggest there have been considerable changes in the structure of insect communities^24,38,39^. Developing comprehensive insect trait datasets, including using proxies and data imputation, will be crucial to address this deficit^40–42^.

In conclusion, despite these reservations, this global analysis suggests that cases of relative increases of larger species^24,43^ or trait constancy despite declines in abundance^44^, may in fact be as frequent as shifts towards smaller sized species^22^. Community responses may therefore be considerably more nuanced and localised than previously considered based on macroecological expectations^8^.

## Methods

### Generating assemblage time series

We downloaded all studies available in the ‘open’ component of the BioTIME database of community time series^14^ from https://doi.org/10.5281/zenodo.3265871. BioTIME contains observations from both fixed plots (repeat measures from the same set of specific localised sites) and from wide ranging surveys and transects that may not necessarily precisely align year-on-year. We followed previous approaches^1^ and first identified studies as ‘multi-site’ or ‘single-site’ based on the number of coordinates in the BioTIME database. Single site studies were considered as one combined assemblage, while widely-dispersed ‘multi-site’ studies were portioned into assemblages based on a global hexagonal grid of 96 km^2^ cells using *dggridR*^45^. We retained records from assemblages with abundance or biomass data of at least 10 distinct species and at least 5 years between the first and the last record.

### Cleaning Names

Although the majority of the records are identified with binomials to species level, a portion of the records in the BioTIME database are labelled only at higher taxonomic levels. For simplicity, we refer to all distinct names as ‘species’. We identified uninformative labels (for example ‘spA’, ‘unidentified’, ‘Miscellaneous’, ‘larvae’, ‘grass’) and common names (mostly birds) were converted to binomials using the Encyclopaedia of Life tool via the *taxize* R package^46,47^ followed by manual inspection based on study location and species distribution where multiple options were presented. We excluded studies where the species are listed using codes. Informative names were standardised against the GBIF name backbone^48^ using *taxize*. The dominant kingdom represented in each study was used to distinguish homonyms. Where BioTIME included only a genus level identification, we matched these to genus level trait values listed in trait databases. Where BioTIME only included taxonomic information of higher rank than genus, we did not attempt to match the traits.

### Trait data

We used four separate trait databases that include some measure of organism size, but we did not mix information between databases. *Amniotes*: The life history database was downloaded from https://doi.org/10.6084/m9.figshare.c.3308127.v1^15^ from which we used the ‘adult_body_mass_g’ field. *Plants*: We downloaded from the TRY database (https://www.try-db.org/)^17^ all records of ‘seed dry mass’ (trait 26) and ‘plant height vegetative’ (trait 3106). We grouped these by accepted species name, and calculated the mean of the log 10 (seedmass) values and the maximum observed height. We did not assign a value when the standard deviation of log 10 (seedmass) values was greater than 1. The resultant dataset was derived from 91 original datasets^51-123^. *Fish*: We downloaded a curated database of fish traits from https://store.pangaea.de/Publications/Beukhof-etal_2019/TraitCollectionFishNAtlanticNEPacificContShelf.xlsx^18^ which in turn is largely based on data from the FishBase database^19^. It is focussed on the North Atlantic and Pacific continental shelf, but this represents the majority of the relevant BioTIME studies. It includes values for both genus and species level. We used maximum length, and when there were multiple values for a particular species, we took an average. *Marine*: We downloaded size data from the World Register of Marine Species (WoRMS) database^16^. Aphia IDs for all the species in our assemblages (excluding plants and fungi) were identified and used to download all attributes associated with these IDs held on WoRMS using the *worrms* R package^49^. Quantitative ‘body size’ measurements of length were scaled to millimetre units. We discarded values from stages other than adults, values corresponding to minimums or thicknesses, then took a mean, except where the values differed by over an order of magnitude, which we discarded. Qualitative body sizes were divided into four categories (<0.2 mm 0.2 2mm, 2-200 mm, >200 mm), that were carried forwards as simple numbers (1-4). Data not from adults was discarded, and where an ID was associated with multiple distinct size categories, it was discarded.

Summaries of the trait data completeness are given in Extended Data Figure 2. Note that 80 studies had sufficient trait data for analysis under multiple traits 36 had both categories of plant data, 31 had length data from both WoRMS and the fish-specific database, 7 studies spanned the amniote life history traits and WoRMS database, 5 studies shared both qualitative and quantitative size information from WoRMS, and one study could be related to both traits from WoRMS and the fish database.

### Abundance change – trait correlation

We assessed each assemblage-trait combination where ≥40% of the species had data for that trait. Within each year, all *n* species in the assemblage were assigned relative ranks (1 = highest, 1/n = lowest) by their abundance or biomass depending on the fields available in BioTIME. Ties were averaged and where a species was not observed in a particular year, it was assigned a rank of zero for that year. Where a study included both ‘abundance’ and ‘biomass’ data, we preferentially used the abundance data. Presence absence-data was not used. We fit a linear model through the relative ranks over time of each species in the assemblage. The set of slopes (β) of these linear models within each dataset summarised the relative change in abundance of each species in the assemblage through time. Very small β values, caused by model fitting errors when there is no change in rank abundance, were set to 0 to avoid spurious rankings. The main response variable ‘τ’ for each assemblage was then computed as Kendall’s rank correlation coefficient between trait values and the set of βs. Species with missing trait values were excluded. The default ‘τ^B^’ approach was used for ties^50^. Where there were multiple assemblages per study, study level τ was a simple arithmetic mean of all assemblage level τ values.

### Statistics

To generate a null model for the impact of traits, the relative abundance rank changes (βs) were computed as above, but the available trait values (including ‘NA’s where trait data was missing) were randomly reassigned to the species in that assemblage and τ recalculated. This was repeated 100 times per assemblage to generate a null distribution of expected τ values for each study. For each tested trait, we used two-sided Welch’s two-sample t-test to determine if the mean τ for that trait was significantly different to this null expectation.

To examine study level determinates of τ within each trait, for each study we calculated: 1) the mean total species richness of each assemblage over the time frame, 2) the mean assemblage level trait data completeness, 3) the mean number of years that from which there was data, 4) the mean span of years from which there was data, 5) the number of assemblages within the study (i.e. the spatial extent), 6) the grain size of the study, as listed in the BioTIME metadata, and 7) the absolute latitude of the centre of the study. We fit a set of linear models to assess whether these factors could predict either τ or τ^2^.

## Author contributions

JCDT designed and conducted the analyses and wrote the first draft of the manuscript. All authors contributed to the manuscript development and revision.

## Acknowledgments

This work would be impossible without all the hundreds of researchers who have contributed to the multiple open-access databases and software used. All authors were supported by NERC grant NE/T003510/1 ‘***Mechanisms and prediction of large-scale ecological responses to environmental change’***.

## Code Availability

All analyses were conducted using R. Code and illustrative notebooks to reproduce all steps is available at: https://github.com/jcdterry/BioTIME_BodySize and archived on Zenodo at https://doi.org/10.5281/zenodo.4745554

## Data Availability

Original sources of open-source datasets are listed in the methods. Processed data are available with analysis code at: https://github.com/jcdterry/BioTIME_BodySize and archived on Zenodo at https://doi.org/10.5281/zenodo.4745554

