## Extended Data Tables and Figures for "No pervasive size trend in global community dynamics"

**Extended Data Table 1** Full statistical results underlying Figure 2. Tests were two-sided Welch's two-sample t-tests, which approximates the relevant degrees of freedom given different variance in the observations and the null model. No corrections were made to the reported values for multiple comparisons.

| Trait | Estimated Difference | Lower CI | Upper CI | Estimated Mean (Observed) | (Null) | t-statistic | p-value | Effective Degrees of Freedom | Sample Size |
|---|---|---|---|---|---|---|---|---|---|
| **Marine** | | | | | | | | | |
| Body Length | 0.0140 | -0.0201 | 0.0481 | 0.0149 | 0.0010 | 0.8173 | 0.4165 | 70.8705 | 71 |
| Qualitative Body Size | 0.0182 | -0.0589 | 0.0954 | 0.0182 | 0.0000 | 0.5422 | 0.6021 | 8.2094 | 9 |
| **Fish** | | | | | | | | | |
| Maximum Length | 0.0030 | -0.0553 | 0.0612 | 0.0007 | -0.0023 | 0.1043 | 0.9175 | 32.2650 | 33 |
| **Amniotes** | | | | | | | | | |
| Adult body mass | 0.0406 | 0.0007 | 0.0806 | 0.0416 | 0.0010 | 2.0446 | **0.0463** | 49.0707 | 49 |
| **Plants** | | | | | | | | | |
| Seed Mass | 0.0005 | -0.0546 | 0.0556 | 0.0021 | 0.0016 | 0.0183 | 0.9855 | 42.6548 | 43 |
| Maximum Height | -0.0103 | -0.0765 | 0.0560 | -0.0095 | 0.0007 | -0.3131 | 0.7559 | 39.5603 | 40 |

**Extended Data Figure 1** Raw relationships between the suite of study-level predictors and the principal response variable $\tau$.

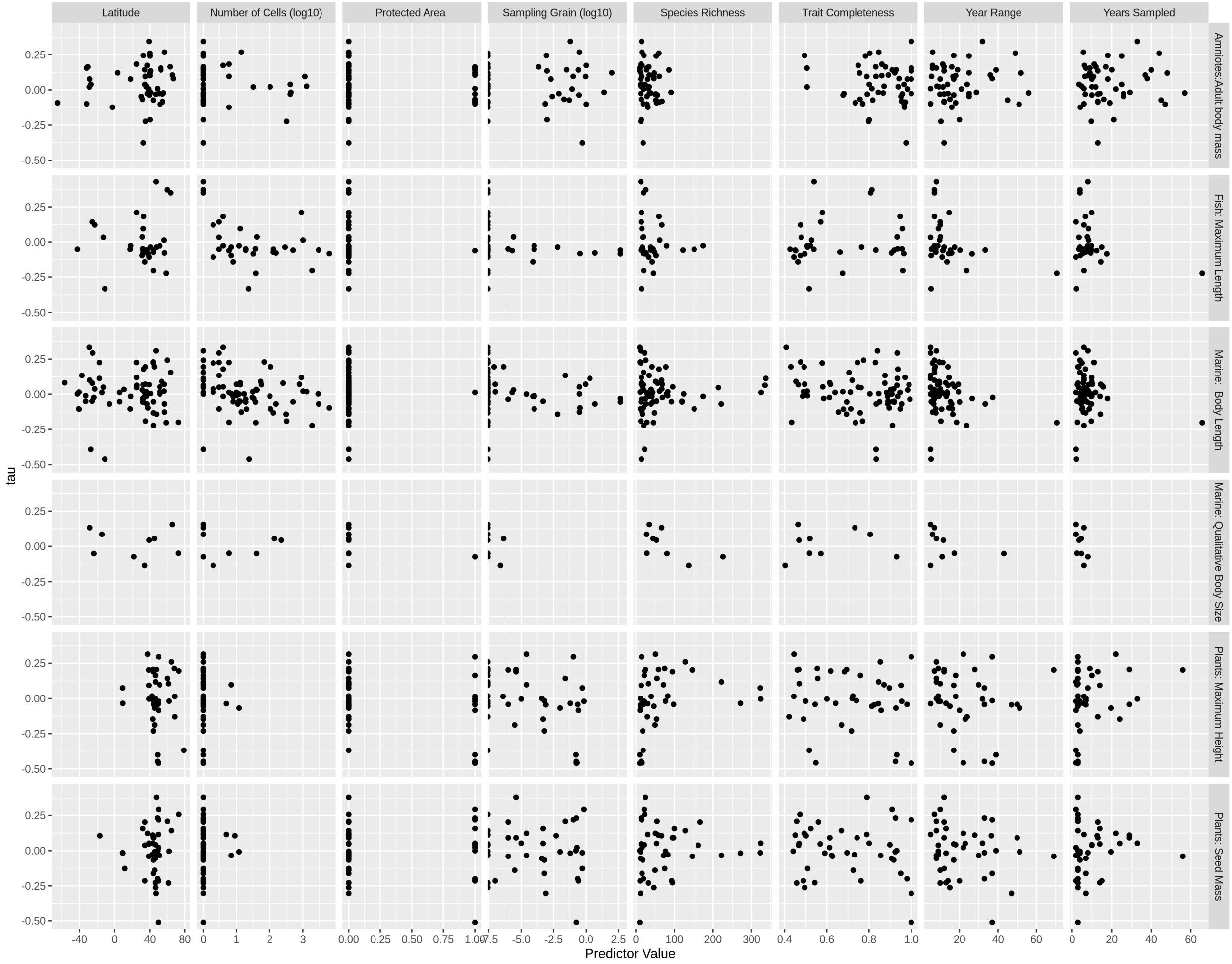

**Extended Data Table 2** Full statistical results of linear models for the putative study-level drivers of $\tau$. Each trait was tested independently and no corrections were made to the reported values for multiple comparisons. There was no consistent drivers - only three relationships were identified as significant at $p<0.05$ (highlighted in **bold**), but these had low explanatory power. Inspection of Extended Data Figure 1 suggests that the impact of number of cells within the fish studies was driven by three outlier single-site studies, while the study-duration and sampling frequency relationships identified for plant height can be seen to be driven by one long term study.

| | Marine | | | | Fish | | Amniotes | | Plants | | | |
|---|---|---|---|---|---|---|---|---|---|---|---|---|
| | Body Length | | Qualitative Body Size | | Maximum Length | | Adult body mass | | Seed Mass | | Maximum Height | |
| | Estimate | p-value | Estimate | p-value | Estimate | p-value | Estimate | p-value | Estimate | p-value | Estimate | p-value |
| **Coefficents** | | | | | | | | | | | | |
| Intercept | 0.2 | 0.0796 | -0.0328 | 0.931 | -0.0578 | 0.635 | 0.16 | 0.32 | -0.0855 | 0.639 | -0.223 | 0.454 |
| Species Richness | -7.86e-05 | 0.758 | -0.0016 | 0.379 | -0.00042 | 0.553 | -0.000755 | 0.518 | 0.000154 | 0.726 | 0.000384 | 0.432 |
| Trait Completeness | -0.147 | 0.176 | 0.233 | 0.57 | 0.112 | 0.431 | -0.157 | 0.355 | 0.0702 | 0.722 | 0.245 | 0.321 |
| Years Sampled | 0.00177 | 0.615 | 0.0144 | 0.778 | -0.00245 | 0.669 | -0.00204 | 0.756 | 0.00543 | 0.17 | 0.0121 | **0.0143** |
| Year Range | -0.00432 | 0.157 | -0.00266 | 0.563 | -0.0021 | 0.691 | 0.00299 | 0.636 | -0.00501 | 0.107 | -0.00782 | **0.0151** |
| Number of Cells (Log10) | -0.0354 | 0.0626 | -0.00322 | 0.96 | -0.0723 | **0.017** | -0.015 | 0.532 | 0.113 | 0.313 | 0.0395 | 0.778 |
| Absolute Latitude | 0.000359 | 0.795 | 0.000226 | 0.946 | 0.00418 | 0.0543 | 0.000679 | 0.728 | 0.0017 | 0.51 | 0.00154 | 0.622 |
| **Summary** | | | | | | | | | | | | |
| Observations | 71 | | 9 | | 33 | | 49 | | 43 | | 40 | |
| Adjusted R-squared | 0.087 | | -0.389 | | 0.261 | | -0.0957 | | -0.025 | | 0.127 | |

**Table S3** Full statistical results of tests for the drivers of $\tau^2$, in order to test if there are drivers for deviations from trait-neutrality. Each trait was tested independently and no corrections were made to the reported values for multiple comparisons. Here also, there were no consistent drivers.

| | Marine | | | | Fish | | Amniotes | | Plants | | | |
|---|---|---|---|---|---|---|---|---|---|---|---|---|
| | Body Length | | Qualitative Body Size | | Maximum Length | | Adult body mass | | Seed Mass | | Maximum Height | |
| | Estimate | p-value | Estimate | p-value | Estimate | p-value | Estimate | p-value | Estimate | p-value | Estimate | p-value |
| **Coefficents** | | | | | | | | | | | | |
| Intercept | 0.0615 | **0.0385** | 0.038 | 0.203 | 0.0357 | 0.32 | 0.0387 | 0.252 | 0.0124 | 0.786 | 0.00756 | 0.932 |
| Species Richness | -0.000192 | **0.00508** | -5.21e-05 | 0.608 | -0.000366 | 0.0852 | -0.000321 | 0.195 | -0.000102 | 0.357 | -0.000116 | 0.429 |
| Trait Completeness | -0.00364 | 0.897 | -0.0355 | 0.232 | 0.000468 | 0.991 | -0.00322 | 0.928 | 0.0161 | 0.745 | -0.00268 | 0.971 |
| Years Sampled | 0.000188 | 0.837 | 0.00182 | 0.572 | -0.000602 | 0.72 | 0.00102 | 0.461 | -0.00131 | 0.189 | -0.00179 | 0.208 |
| Year Range | -0.000125 | 0.874 | -0.00013 | 0.634 | 0.000635 | 0.681 | -0.000819 | 0.538 | 0.000768 | 0.32 | 0.00181 | 0.056 |
| Number of Cells (Log10) | -0.00953 | 0.054 | -0.00848 | 0.134 | -0.0218 | **0.014** | -0.0033 | 0.514 | -0.0427 | 0.132 | -0.0647 | 0.128 |
| Absolute Latitude | -0.000406 | 0.26 | -7.5e-05 | 0.714 | 0.000908 | 0.146 | -0.000156 | 0.703 | 0.000341 | 0.599 | 0.000488 | 0.602 |
| **Summary** | | | | | | | | | | | | |
| Observations | 71 | | 9 | | 33 | | 49 | | 43 | | 40 | |
| Adjusted R-squared | 0.0692 | | 0.3 | | 0.182 | | -0.0531 | | 0.0922 | | 0.0884 | |

**Extended Data Figure 2** Further details of degree of overlap and correspondence between traits. a) Number of species that could be related to at least one trait from the four sources. b) Overlap within the WoRMS database between the quantitative and qualitative body lengths was relatively low. In cases where the data was available on both categories, the Spearman's rank correlation was 0.65. c) Very strong correlation between the size traits for species that had data in both the WoRMS and the FishBase databases. d) Overlap in trait data between then plant species held in the TRY database was comparatively high. e) Correlation between the seed mass and vegetative height trait values was moderate, and considerably less within guilds.

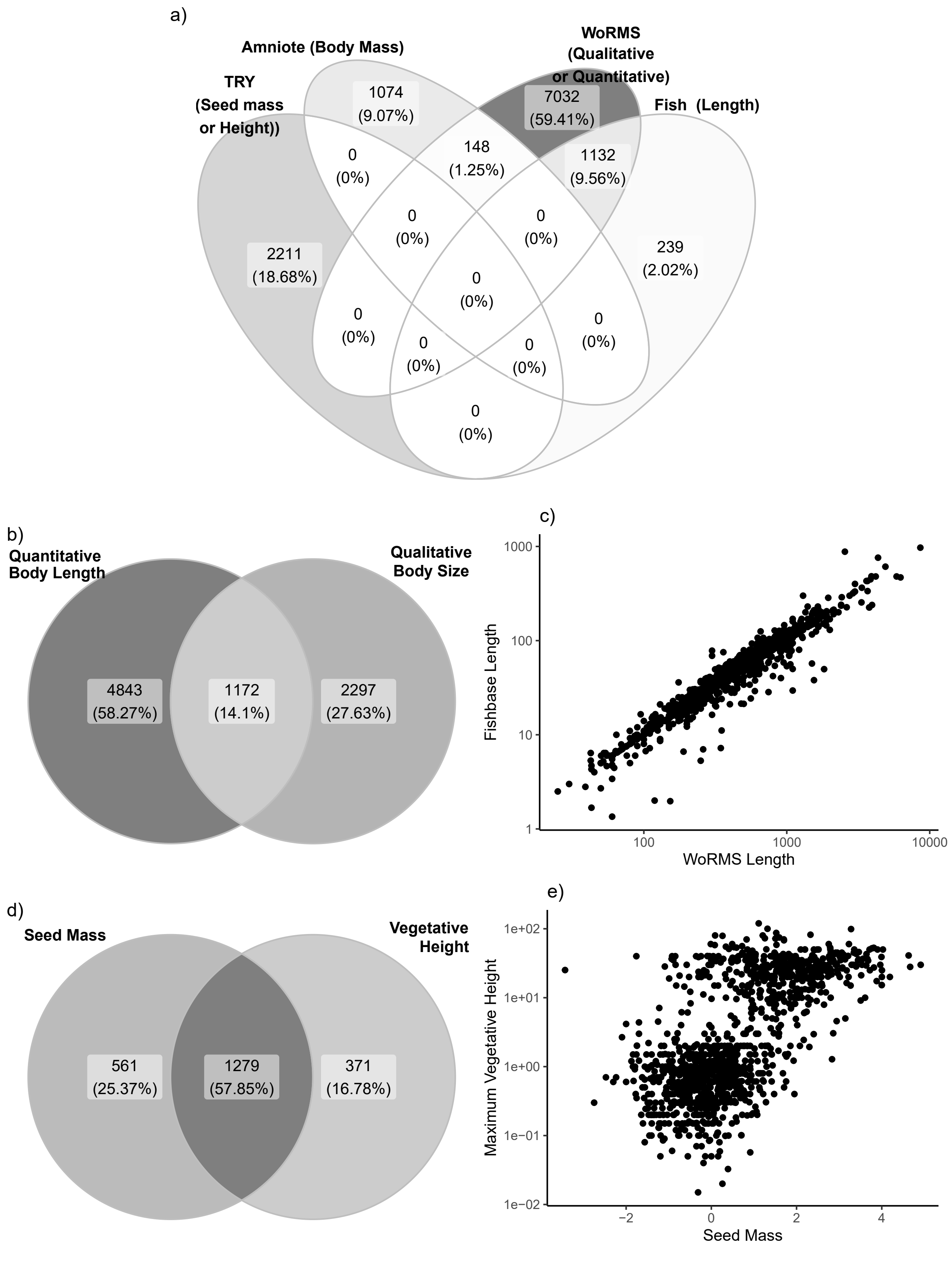
a)
Amniote (Body Mass)
WoRMS
(Qualitative
or Quantitative)
TRY
(Seed mass
or Height))
Fish (Length)
1074
(9.07%)
7032
(59.41%)
148
(1.25%)
1132
(9.56%)
2211
(18.68%)
239
(2.02%)
0
(0%)
b)
Quantitative
Body Length
Qualitative
Body Size
4843
(58.27%)
1172
(14.1%)
2297
(27.63%)
c)
Fishbase Length
WoRMS Length
1000
100
10
1
100
1000
10000
d)
Seed Mass
Vegetative
Height
561
(25.37%)
1279
(57.85%)
371
(16.78%)
e)
Maximum Vegetative Height
Seed Mass
1e+02
1e+01
1e+00
1e−01
1e−02
−2
0
2
4

**Extended Data Table 4 -** List of studies used from BioTIME, including key statistics and description. Studies are ordered by trait, then by $\tau$. If a study could be linked to multiple traits, it will therefore appear multiple times. Note this table is also available in .csv format. ID numbers correspond to the BioTIME database, and columns ‘Latitude’ to ‘Data Source’ are directly drawn from the BioTIME metadata.

| Trait tested | ID | tau | N. Sp. | N. Years | Trait % | Year Span | N. Cells | Latitude | Longitude | Grain | Abundance | Biomass | Study Title | Data Source |
|---|---|---|---|---|---|---|---|---|---|---|---|---|---|---|
| Amniotes:Adult body mass | 321 | -0.376 | 19.0 | 13.000 | 98 | 12.0 | 1 | 32.550 | -106.812 | 0.500 | Count | Weight | Small Mammal Exclosure Study. Jornada LTER. SMES rodent trapping data | http://jornada.nmsu.edu/lter/dataset/497[illegible] |
| Amniotes:Adult body mass | 172 | -0.224 | 13.0 | 9.696 | 80 | 10.4 | 326 | 35.010 | -24.225 | 0.000 | Count | NA | POPA cetacean. seabird. and sea turtle sightings in the Azores area 1998-2009 (OBIS SEAMAP) | http://www.iobis.org/mapper/?dataset=4[illegible] |
| Amniotes:Adult body mass | 308 | -0.212 | 14.0 | 21.000 | 80 | 20.0 | 1 | 40.171 | -79.260 | 0.001 | Count | Weight | Powdermill Nature Reserve monitored small mammal populations from 1979-1999. | https://portal.lternet.edu/nis/metadatavi[illegible] lter-vcr.67.17 |
| Amniotes:Adult body mass | 516 | -0.123 | 31.5 | 4.167 | 97 | 16.0 | 6 | -2.386 | -59.919 | 0.000 | Count | NA | A large-scale fragmentation experiment for Neotropical bats | None |
| Amniotes:Adult body mass | 46 | -0.102 | 29.0 | 47.000 | 97 | 51.0 | 1 | 51.698 | -5.277 | 1.000 | Count | NA | Skokholm Bird Observatory | http://ecologicaldata.org/wiki/skokholm-bird-observatory |
| Amniotes:Adult body mass | 275 | -0.099 | 19.0 | 6.000 | 77 | 5.0 | 1 | -31.968 | 115.833 | 0.001 | Count | Size | Response of an urban remnant reptile community to summer wildfire | http://journals.plos.org/plosone/article?i[illegible] |
| Amniotes:Adult body mass | 419 | -0.092 | 54.0 | 19.000 | 73 | 18.0 | 1 | -64.770 | -64.050 | 0.000 | Count | NA | Data collected aboard cruises off the coast of the Western Antarctic Penninsula | https://portal.lternet.edu/nis/mapbrowse[illegible] lter-pal.100.1 |
| Amniotes:Adult body mass | 441 | -0.087 | 61.0 | 13.000 | 97 | 12.0 | 1 | 54.504 | 60.294 | 0.000 | Count | NA | Long-term dynamics of bird populations in birch forests of Ilmen Nature Reserve during the breeding period individuals / km2 | http://ashipunov.info/shipunov/school/b[illegible] |
| Amniotes:Adult body mass | 440 | -0.081 | 68.0 | 13.000 | 95 | 12.0 | 1 | 54.504 | 60.294 | 0.000 | Density | NA | Long-term dynamics of bird populations in pine-birch forests of Ilmen Nature Reserve during the breeding period individuals / km2 | http://ashipunov.info/shipunov/school/b[illegible] |
| Amniotes:Adult body mass | 39 | -0.073 | 52.0 | 45.000 | 82 | 45.0 | 1 | 43.910 | -71.750 | 0.050 | Density | NA | Bird community dynamics in a temperate deciduous forest Long-term trends at Hubbard Brook | http://www.esajournals.org/toc/emon/56[illegible] |
| Amniotes:Adult body mass | 358 | -0.067 | 14.0 | 16.000 | 76 | 15.0 | 1 | 31.583 | -94.817 | 0.020 | Density | NA | Neotropical Migratory Bird Communities in a Developing Pine Plantation | http://www.srs.fs.fed.us/pubs/520 |
| Amniotes:Adult body mass | 59 | -0.046 | 29.0 | 26.000 | 95 | 25.0 | 1 | 30.323 | -103.501 | 0.002 | Count | NA | Long-term monitoring and experimental manipulation of a Chihuahuan Desert ecosystem near Portal. Arizona. USA | http://esapubs.org/archive/ecol/E090/11[illegible] |
| Amniotes:Adult body mass | 41 | -0.036 | 56.0 | 10.000 | 68 | 17.0 | 1 | 39.500 | -82.480 | 0.280 | Count | NA | Time and space and the variation of species | http://www.esajournals.org/toc/ecol/41/[illegible] |
| Amniotes:Adult body mass | 217 | -0.031 | 50.5 | 6.664 | 79 | 9.9 | 420 | 46.829 | -109.982 | 0.000 | Count | NA | Landbird Monitoring Program (UMT-LBMP) | http://www.treesearch.fs.fed.us/pubs/320[illegible] |
| Amniotes:Adult body mass | 439 | -0.029 | 52.0 | 13.000 | 96 | 12.0 | 1 | 54.504 | 60.294 | 0.000 | Density | NA | Long-term dynamics of bird populations in pine forests of Ilmen Nature Reserve during the breeding period individuals / km2 | http://ashipunov.info/shipunov/school/b[illegible] |

*(continued)*

| Trait tested | ID | tau | N. Sp. | N. Years | Trait % | Year Span | N. Cells | Latitude | Longitude | Grain | Abundance | Biomass | Study Title | Data Source |
|---|---|---|---|---|---|---|---|---|---|---|---|---|---|---|
| Amniotes:Adult body mass | 319 | -0.026 | 35.0 | 14.000 | 68 | 14.0 | 1 | 37.250 | -96.717 | 0.008 | Count | NA | Effects of rangeland management on community dynamics of herpetofauna to the tallgrass prairie | http://www.bioone.org/doi/pdf/10.1655/ 0831%282006%2962%5B378%3AEORMO |
| Amniotes:Adult body mass | 47 | -0.026 | 13.0 | 26.000 | 100 | 25.0 | 1 | 50.845 | -107.446 | 0.000 | Count | NA | Detection of Density-Dependent Effects in Annual Duck Censuses | http://www.esajournals.org/toc/ecol/65/ |
| Amniotes:Adult body mass | 339 | -0.022 | 39.0 | 57.000 | 87 | 56.0 | 1 | 55.717 | 13.333 | 0.000 | Count | NA | Species trends turnover and composition of a woodland bird community in southern Sweden during a period of 57 years. | http://springer |
| Amniotes:Adult body mass | 195 | -0.017 | 91.5 | 29.305 | 84 | 28.8 | 439 | 40.809 | -96.187 | 25.427 | Count | NA | Breeding birds survey North America | https://www.pwrc.usgs.gov/bbs/ |
| Amniotes:Adult body mass | 361 | 0.006 | 20.0 | 22.000 | 98 | 21.0 | 1 | 38.610 | -79.835 | 0.083 | Count | NA | A long-term bird population study in an Appalachian spruce forest | http://www.jstor.org/stable/4161914. |
| Amniotes:Adult body mass | 442 | 0.009 | 33.0 | 6.000 | 81 | 5.0 | 1 | 48.669 | 85.654 | 0.000 | Count | NA | Composition and abundance of bird species in the village Matabay in June 1980-1985 (absolute indicators (area 025 km2)) | http://cyberleninka.ru/article/n/ptitsy-naselyonnyh-punktov-markakolskoy-kotloviny-yuzhnyy-altay |
| Amniotes:Adult body mass | 67 | 0.021 | 15.4 | 11.906 | 51 | 11.4 | 32 | -28.954 | 24.951 | 0.000 | Count | NA | Animal Demography Unit - Coordinated Waterbird Counts (CWAC) (AfrOBIS) | http://www.iobis.org/mapper/?dataset= |
| Amniotes:Adult body mass | 374 | 0.022 | 34.9 | 10.067 | 94 | 9.4 | 104 | 35.961 | 136.046 | 0.000 | Count | NA | Monitoring site 1000 Shorebird Survey | http://www.biodic.go.jp/moni1000/findin |
| Amniotes:Adult body mass | 166 | 0.026 | 13.0 | 4.926 | 87 | 8.3 | 1298 | 36.075 | -70.992 | 0.000 | Count | NA | PIROP Northwest Atlantic 1965-1992 (SEAMAP) | http://www.iobis.org/mapper/?dataset= |
| Amniotes:Adult body mass | 366 | 0.039 | 24.0 | 25.000 | 97 | 24.0 | 1 | 34.350 | -106.880 | 0.000 | Count | NA | Small Mammal Exclosure Study (SMES) | https://portal.lternet.edu/nis/mapbrowse lter-sev.8.297976 |
| Amniotes:Adult body mass | 108 | 0.039 | 12.2 | 3.524 | 80 | 13.7 | 420 | -27.174 | 3.946 | 0.000 | Count | NA | Seabirds of the Southern and South Indian Ocean (Australian Antarctic Data Centre) | https://data.aad.gov.au/ |
| Amniotes:Adult body mass | 58 | 0.077 | 31.0 | 18.000 | 98 | 17.0 | 1 | 18.190 | -65.430 | 0.002 | Count | NA | Avian populations long-term monitoring dataset. San Juan. Puerto Rico Luquillo Long Term Ecological Research Site Database Grid points bird counts DBAS 23 | http://luq.lternet.edu/data/luqmetadata |
| Amniotes:Adult body mass | 348 | 0.077 | 13.0 | 10.000 | 100 | 10.0 | 1 | -28.609 | -48.981 | 0.000 | Count | Weight | Bats (Mammalia Chiroptera) in restinga in the municipality of Jaguaruna south of Santa Catarina Brazil. | None |
| Amniotes:Adult body mass | 420 | 0.080 | 47.0 | 38.000 | 94 | 37.0 | 1 | 67.077 | 17.435 | 0.000 | Count | NA | Species composition and population fluctuations of alpine bird communities during 38 years in the Scandinavian mountain range | http://www.luvre.org/data_o_pdf/Luvre 135%202006%20Svensson%20Heden%2038 |

*(continued)*

| Trait tested | ID | tau | N. Sp. | N. Years | Trait % | Year Span | N. Cells | Latitude | Longitude | Grain | Abundance | Biomass | Study Title | Data Source |
|---|---|---|---|---|---|---|---|---|---|---|---|---|---|---|
| Amniotes:Adult body mass | 169 | 0.095 | 15.4 | 10.799 | 79 | 16.7 | 1152 | 34.858 | -121.615 | 0.900 | Count | NA | CalCOFI and NMFS Seabird and Marine Mammal Observation Data. 1987-2006 (SEAMAP) | http://www.iobis.org/mapper/?dataset=2 |
| Amniotes:Adult body mass | 372 | 0.096 | 61.0 | 6.500 | 83 | 5.5 | 6 | 34.952 | 134.975 | 0.100 | Count | NA | Monitoring site 1000 Village survey - Bird survey data | http://www.biodic.go.jp/moni1000/findin |
| Amniotes:Adult body mass | 415 | 0.101 | 45.0 | 8.000 | 86 | 18.0 | 1 | 40.133 | -88.300 | 0.000 | Density | NA | Bird populations in east central Illinois. Fluctuations variations and development over a half-century | https://www.ideals.illinois.edu/handle/21 |
| Amniotes:Adult body mass | 363 | 0.105 | 35.0 | 37.000 | 89 | 36.0 | 1 | 65.968 | 16.317 | 0.000 | Count | NA | The 37-year dynamics of a subalpine bird community with special emphasis on the influence of environmental temperature and Epirrita autumnata cycles. | http://www.luvre.org/data_o_pdf/Luvre 130%202004%20Enemar%20mfl%2037%20 |
| Amniotes:Adult body mass | 414 | 0.120 | 48.0 | 48.000 | 75 | 52.0 | 1 | 39.983 | -88.650 | 0.000 | Density | NA | Bird populations in east central Illinois. Fluctuations variations and development over a half-century | https://www.ideals.illinois.edu/handle/21 |
| Amniotes:Adult body mass | 312 | 0.121 | 13.0 | 9.000 | 92 | 25.0 | 1 | 3.500 | 35.750 | 100.000 | Count | NA | Stability in a Multi-Species Assemblage of Large Herbivores in East Africa | http://www.jstor.org/stable/4219351?seq |
| Amniotes:Adult body mass | 357 | 0.135 | 10.0 | 13.000 | 100 | 12.0 | 1 | 40.829 | -104.758 | 0.001 | Count | NA | Small Mammal Trapping Webs on the Central Plains Experimental Range | https://portal.lternet.edu/nis/mapbrowse lter-sgs.137.17 |
| Amniotes:Adult body mass | 360 | 0.142 | 86.0 | 40.000 | 91 | 39.0 | 1 | 52.717 | 24.267 | 0.250 | NA | Weight | Bialowieza National Park bird assemblage | http://www.bioone.org/doi/abs/10.3161/ |
| Amniotes:Adult body mass | 56 | 0.143 | 28.0 | 20.000 | 93 | 19.0 | 1 | 34.200 | -106.430 | 0.031 | Count | Weight | Small Mammal Mark-Recapture Population Dynamics at Core Research Sites | http://sev.lternet.edu/data/sev-8 |
| Amniotes:Adult body mass | 381 | 0.155 | 29.0 | 7.000 | 51 | 6.0 | 1 | -31.933 | 115.767 | 0.000 | Count | NA | Long-term sampling of a herpetofaunal assemblage on an isolated urban bushland | http://www.rswa.org.au/publications/Jou |
| Amniotes:Adult body mass | 447 | 0.156 | 10.0 | 9.000 | 100 | 8.0 | 1 | 52.601 | 38.928 | 0.000 | Density | NA | Long-term population dynamics of small mammals in the Natural Boundary Morozova Gora (individuals / 100 trap-nights) | http://elibrary.ru/item.asp?id=24990048 |
| Amniotes:Adult body mass | 327 | 0.164 | 12.0 | 17.000 | 88 | 16.0 | 1 | -30.600 | -71.700 | 0.000 | Count | Weight | Fray Jorge Small Mammals 1989-2005 | http://esapubs.org/archive/ecol/E094/08 |
| Amniotes:Adult body mass | 475 | 0.164 | 34.0 | 12.000 | 83 | 12.0 | 1 | 63.417 | 10.500 | 0.000 | Count | NA | Structure and dynamics of a passerine bird community in a spruce-dominated boreal forest | http://www.sekj.org/PDF/anzf30/anz30-043-054.pdf |
| Amniotes:Adult body mass | 373 | 0.174 | 16.0 | 6.250 | 75 | 6.0 | 4 | 37.071 | 137.152 | 1.000 | Count | NA | Village survey Medium and large mammal survey data | http://www.biodic.go.jp/moni1000/findin |

*(continued)*

| Trait tested | ID | tau | N. Sp. | N. Years | Trait % | Year Span | N. Cells | Latitude | Longitude | Grain | Abundance | Biomass | Study Title | Data Source |
|---|---|---|---|---|---|---|---|---|---|---|---|---|---|---|
| Amniotes:Adult body mass | 171 | 0.183 | 13.3 | 11.167 | 86 | 11.3 | 6 | 24.896 | -76.292 | 0.000 | Count | NA | Bahamas Marine Mammal Research Organisation Opportunistic Sightings (SEAMAP) | http://www.iobis.org/mapper/?dataset=2 |
| Amniotes:Adult body mass | 416 | 0.241 | 53.0 | 25.000 | 78 | 25.0 | 1 | 40.133 | -88.300 | 0.000 | Density | NA | Bird populations in east central Illinois. Fluctuations variations and development over a half-century | https://www.ideals.illinois.edu/handle/21 |
| Amniotes:Adult body mass | 316 | 0.244 | 21.0 | 18.000 | 49 | 17.0 | 1 | 32.620 | -106.740 | 0.001 | Count | Weight | Lizard pitfall trap data (LTER-II LTER-III) | http://jornada.nmsu.edu/lter/dataset/498 |
| Amniotes:Adult body mass | 413 | 0.260 | 60.0 | 44.000 | 80 | 49.0 | 1 | 39.983 | -88.650 | 0.000 | Density | NA | Bird populations in east central Illinois. Fluctuations variations and development over a half-century | https://www.ideals.illinois.edu/handle/21 |
| Amniotes:Adult body mass | 91 | 0.268 | 15.9 | 5.714 | 85 | 6.1 | 14 | 57.039 | 20.556 | 0.300 | Count | NA | Baltic seabirds transect surveys | http://www.iobis.org/mapper/?dataset=2 |
| Amniotes:Adult body mass | 311 | 0.345 | 15.0 | 33.000 | 100 | 32.0 | 1 | 39.083 | -96.583 | 0.060 | Count | NA | Seasonal summary of numbers of small mammals on 14 LTER traplines in prairie habitats at Konza Prairie | http://lter.konza.ksu.edu/content/csm01-seasonal-summary-numbers-small-mammals-14-lter-traplines-prairie-habitats-konza |
| Marine: Body Length | 284 | -0.461 | 14.5 | 2.208 | 83 | 5.2 | 24 | -11.206 | -43.617 | 0.000 | Count | Weight | Previous__fisheries__REVIZEE_Program (Tropical and Subtropical Western South Atlantic OBIS) | http://www.iobis.org/mapper/?dataset=2 |
| Marine: Body Length | 142 | -0.392 | 23.0 | 2.000 | 83 | 5.0 | 1 | -27.328 | -45.330 | 0.000 | Count | NA | Pelagic and Demersal Fish Database II. REVIZEE South Area (WSAOBIS) | http://www.iobis.org/mapper/?dataset=1 |
| Marine: Body Length | 119 | -0.223 | 20.5 | 6.017 | 91 | 23.7 | 1896 | 43.987 | -63.670 | 0.000 | Count | Weight | DFO Maritimes Research Vessel Trawl Surveys Fish Observations (OBIS Canada) | http://www.iobis.org/mapper/?dataset=2 |
| Marine: Body Length | 428 | -0.202 | 46.0 | 65.737 | 74 | 70.6 | 38 | 58.959 | 9.768 | 0.000 | Count | NA | Long term monitoring of fish abundances from coastal SKagerrak | None |
| Marine: Body Length | 97 | -0.200 | 28.8 | 2.750 | 43 | 18.5 | 6 | 72.739 | 10.694 | 0.000 | Count | NA | Archives of the Arctic Seas Zooplankton (ARC) | http://www.iobis.org/mapper/?dataset=4 |
| Marine: Body Length | 172 | -0.191 | 13.0 | 9.696 | 77 | 10.4 | 326 | 35.010 | -24.225 | 0.000 | Count | NA | POPA cetacean. seabird. and sea turtle sightings in the Azores area 1998-2009 (OBIS SEAMAP) | http://www.iobis.org/mapper/?dataset=4 |
| Marine: Body Length | 182 | -0.142 | 16.9 | 14.321 | 69 | 16.6 | 315 | 47.481 | -62.762 | 0.006 | Count | NA | Snow crab research trawl survey database (Southern Gulf of St. Lawrence. Gulf region. Canada) from 1988 to 2010 (OBIS Canada) | http://www.iobis.org/mapper/?dataset=2 |
| Marine: Body Length | 123 | -0.132 | 48.6 | 6.885 | 76 | 7.7 | 130 | 43.891 | -68.941 | 0.000 | Count | Weight | Maine Department of Marine Resources Inshore Trawl Survey 2000?2009 (OBIS-USA) | http://www.iobis.org/mapper/?dataset=1 |
| Marine: Body Length | 91 | -0.127 | 15.9 | 5.714 | 65 | 6.1 | 14 | 57.039 | 20.556 | 0.300 | Count | NA | Baltic seabirds transect surveys | http://www.iobis.org/mapper/?dataset=2 |

*(continued)*

| Trait tested | ID | tau | N. Sp. | N. Years | Trait % | Year Span | N. Cells | Latitude | Longitude | Grain | Abundance | Biomass | Study Title | Data Source |
|---|---|---|---|---|---|---|---|---|---|---|---|---|---|---|
| Marine: Body Length | 432 | -0.106 | 12.7 | 5.000 | 68 | 10.8 | 20 | -40.493 | 172.716 | 0.000 | Count | NA | The New Zealand Freshwater Fish Database - Observation (Spotlighting visual) | https://www.niwa.co.nz/our-services/online-services/freshwater-fish-database |
| Marine: Body Length | 190 | -0.104 | 151.3 | 8.667 | 95 | 7.7 | 3 | 17.757 | -64.604 | 0.000 | Count | NA | St. Croix. USVI Fish Assessment and Monitoring Data (2002 - Present) (NOAA-CCMA) | http://www.iobis.org/mapper/?dataset=1 |
| Marine: Body Length | 430 | -0.104 | 11.9 | 5.189 | 71 | 16.3 | 106 | -40.943 | 172.499 | 0.000 | Count | NA | The New Zealand Freshwater Fish Database - Electric fishing - Backpack | https://www.niwa.co.nz/our-services/online-services/freshwater-fish-database |
| Marine: Body Length | 180 | -0.097 | 19.4 | 5.252 | 90 | 14.4 | 6328 | 37.771 | -50.793 | 0.333 | Count | NA | ECNASAP - East Coast North America Strategic Assessment (OBIS Canada) | http://www.iobis.org/mapper/?dataset=3 |
| Marine: Body Length | 436 | -0.069 | 221.4 | 4.636 | 95 | 6.6 | 11 | -5.819 | 110.363 | 0.000 | Density | Weight | Karimunjawa WCS fish survey | None |
| Marine: Body Length | 213 | -0.069 | 39.6 | 9.802 | 83 | 33.4 | 2996 | 36.625 | -72.636 | 0.000 | Count | Weight | Northeast Fisheries Science Center Bottom Trawl Survey Data (OBIS-USA) | http://www.iobis.org/mapper/?dataset=1 |
| Marine: Body Length | 466 | -0.069 | 26.7 | 9.410 | 89 | 15.9 | 156 | 56.991 | -9.073 | 5.000 | Count | NA | Trawl Survey Data from Rockall Scotland (1986 - 2008) | http://www.gov.scot/Topics/marine/scier |
| Marine: Body Length | 212 | -0.056 | 44.6 | 4.162 | 92 | 5.6 | 37 | 31.941 | -78.825 | 0.000 | Count | Weight | MARMAP Yankee Trawl 1990-2009 | http://www.usgs.gov/obis-usa/search/?datasetid=MARMAP__Yanke |
| Marine: Body Length | 505 | -0.055 | 119.6 | 7.632 | 69 | 20.1 | 19 | 32.133 | 34.668 | 445.000 | Count | NA | Fish and marine invertebrates from the Israeli Eastern Mediterranean sea 1990-4, 2000, 2008-2012 | none |
| Marine: Body Length | 288 | -0.054 | 13.6 | 5.262 | 85 | 15.3 | 511 | 43.977 | -63.682 | 0.000 | Count | NA | DFO Maritimes Research Vessel Trawl Surveys Fish Observations (OBIS Canada) | http://www.iobis.org/mapper/?dataset=2 |
| Marine: Body Length | 437 | -0.053 | 92.4 | 3.375 | 91 | 5.4 | 8 | 5.753 | 95.280 | 0.000 | Density | Weight | Aceh WCS fish survey 2010 to 2016 | None |
| Marine: Body Length | 295 | -0.050 | 119.5 | 6.000 | 92 | 6.4 | 13 | -33.051 | 139.184 | 0.000 | Count | Weight | Systematic global assessment of reef fish communities by the Reef Life Survey program | http://reeflifesurvey.imas.utas.edu.au/por |
| Marine: Body Length | 290 | -0.049 | 53.9 | 2.500 | 89 | 6.2 | 12 | -25.813 | 135.278 | 0.000 | Count | Weight | CSIRO Marine Data Warehouse (OBIS Australia) | http://www.iobis.org/mapper/?dataset=5 |
| Marine: Body Length | 127 | -0.036 | 14.7 | 6.421 | 99 | 6.8 | 19 | 32.335 | -78.992 | 0.000 | Count | Weight | MARMAP Florida Antillean Trap Survey 1990-2009 (OBIS-USA) | http://www.iobis.org/mapper/?dataset=1 |
| Marine: Body Length | 507 | -0.031 | 25.7 | 17.833 | 59 | 26.7 | 33 | 31.912 | 34.392 | 445.000 | NA | Weight | Trawl fisheries in Israeli Mediterranean | none |
| Marine: Body Length | 330 | -0.023 | 69.1 | 4.250 | 61 | 37.2 | 30 | -23.831 | 136.449 | 0.000 | Density | NA | Over 75 years of zooplankton data from Australia | http://esapubs.org/archive/ecol/E095/27 |
| Marine: Body Length | 271 | -0.016 | 41.8 | 14.125 | 89 | 13.1 | 8 | 34.306 | -119.875 | 0.000 | Count | NA | Santa Barbara Coastal LTER | http://sbc.lternet.edu//index.html |
| Marine: Body Length | 189 | -0.016 | 175.0 | 9.750 | 94 | 8.8 | 4 | 18.309 | -64.749 | 0.000 | Count | NA | St. John. USVI Fish Assessment and Monitoring Data (2002 - Present) (NOAA-CCMA) | http://www.iobis.org/mapper/?dataset=1 |
| Marine: Body Length | 374 | -0.015 | 35.7 | 9.944 | 49 | 9.3 | 102 | 35.961 | 136.046 | 0.000 | Count | NA | Monitoring site 1000 Shorebird Survey | http://www.biodic.go.jp/moni1000/findin |

*(continued)*

| Trait tested | ID | tau | N. Sp. | N. Years | Trait % | Year Span | N. Cells | Latitude | Longitude | Grain | Abundance | Biomass | Study Title | Data Source |
|---|---|---|---|---|---|---|---|---|---|---|---|---|---|---|
| Marine: Body Length | 296 | -0.011 | 82.7 | 5.800 | 51 | 6.3 | 10 | -33.051 | 139.184 | 0.000 | Count | NA | Systematic global assessment of reef fish communities by the Reef Life Survey program | http://reeflifesurvey.imas.utas.edu.au/por |
| Marine: Body Length | 371 | 0.000 | 32.0 | 6.000 | 50 | 5.0 | 1 | 36.705 | 137.265 | 0.000 | NA | Cover | Monitoring site 1000 coastal zone research - Algae survey transects | http://www.biodic.go.jp/moni1000/findin |
| Marine: Body Length | 501 | 0.000 | 79.8 | 7.917 | 91 | 6.9 | 12 | 51.530 | 2.730 | 0.000 | Count | NA | Epibenthos and demersal fish monitoring at long-term monitoring stations in the Belgian part of the North Sea | none |
| Marine: Body Length | 500 | 0.001 | 124.1 | 9.588 | 50 | 9.1 | 17 | 51.495 | 2.812 | 0.300 | Density | NA | Macrobenthos monitoring at long-term monitoring stations in the Belgian part of the North Sea from 2001 on | none |
| Marine: Body Length | 148 | 0.002 | 29.3 | 4.147 | 80 | 13.6 | 2898 | -42.407 | 172.260 | 0.000 | NA | Weight | South Western Pacific Regional OBIS Data provider for the NIWA Marine Biodata Information System (South Western Pacific OBIS) | http://www.gbif.org/dataset/83b58bb2-f762-11e1-a439-00145eb45e9a |
| Marine: Body Length | 467 | 0.004 | 38.6 | 4.000 | 85 | 12.0 | 7 | 40.676 | -8.685 | 0.000 | Count | Weight | Spatial and temporal organisation of a coastal lagoon fish community | None |
| Marine: Body Length | 133 | 0.012 | 28.0 | 2.000 | 64 | 6.0 | 1 | -14.617 | -42.219 | 0.000 | Count | NA | Copepods (Tropical and Subtropical Western South Atlantic OBIS) | http://www.iobis.org/mapper/?dataset=5 |
| Marine: Body Length | 359 | 0.012 | 43.8 | 11.833 | 88 | 10.8 | 6 | 34.309 | -119.874 | 0.000 | Count | NA | SBC LTER Reef Kelp Forest Community Dynamics Fish abundance | https://portal.lternet.edu/nis/mapbrowse lter-sbc.17.27 |
| Marine: Body Length | 438 | 0.012 | 325.0 | 6.000 | 95 | 7.1 | 10 | 5.765 | 95.219 | 0.000 | Density | Weight | Aceh WCS fish surveys | None |
| Marine: Body Length | 431 | 0.014 | 11.0 | 5.500 | 71 | 13.5 | 2 | -40.634 | 172.450 | 0.000 | Count | NA | The New Zealand Freshwater Fish Database - Traps- Gee Minnow traps | https://www.niwa.co.nz/our-services/online-services/freshwater-fish-database |
| Marine: Body Length | 204 | 0.016 | 82.9 | 5.516 | 49 | 19.5 | 31 | 51.439 | 2.683 | 0.000 | Count | NA | MACROBEL Long term trends in the macrobenthos of the Belgian Continental Shelf | http://www.emodnet-biology.eu/data-catalog?module=dataset&dasid=145 |
| Marine: Body Length | 166 | 0.017 | 13.0 | 4.928 | 68 | 8.3 | 1298 | 36.075 | -70.992 | 0.000 | Count | NA | PIROP Northwest Atlantic 1965-1992 (SEAMAP) | http://www.iobis.org/mapper/?dataset=2 |
| Marine: Body Length | 163 | 0.021 | 61.6 | 8.261 | 51 | 9.7 | 1015 | 56.500 | -168.150 | 0.000 | Count | Weight | North Pacific Groundfish Observer (North Pacific Research Board) | http://www.iobis.org/mapper/?dataset=6 |
| Marine: Body Length | 125 | 0.029 | 19.7 | 7.707 | 98 | 10.1 | 41 | 31.442 | -78.849 | 0.000 | Count | Weight | MARMAP Chevron Trap Survey 1990-2009 (OBIS-USA) | http://www.iobis.org/mapper/?dataset=1 |
| Marine: Body Length | 121 | 0.033 | 39.8 | 4.846 | 90 | 7.0 | 39 | 10.778 | -169.454 | 0.000 | Count | NA | CRED Rapid Ecological Assessments of Fish Belt Transect Surveys and Fish Stationary Point Count Surveys in the Pacific Ocean 2000-2010 (OBIS-USA) | http://www.iobis.org/mapper/?dataset=1 |
| Marine: Body Length | 511 | 0.037 | 67.5 | 6.000 | 92 | 10.0 | 2 | -22.821 | -43.149 | 0.000 | Count | Weight | Demersal fish hauls from Guanabara Bay Brazil 2005-2015 | None |

*(continued)*

| Trait tested | ID | tau | N. Sp. | N. Years | Trait % | Year Span | N. Cells | Latitude | Longitude | Grain | Abundance | Biomass | Study Title | Data Source |
|---|---|---|---|---|---|---|---|---|---|---|---|---|---|---|
| Marine: Body Length | 412 | 0.046 | 214.0 | 6.000 | 76 | 5.0 | 1 | 25.042 | 121.942 | 0.000 | Density | NA | Icthyoplankton data collected from Yenliao Bay in 6 stations northeast of Taiwan (1995-2000) | None |
| Marine: Body Length | 135 | 0.049 | 18.0 | 3.333 | 75 | 5.0 | 3 | -13.022 | -45.166 | 0.000 | NA | Weight | Previous_fisheries_REVIZEE_Program (Tropical and Subtropical Western South Atlantic OBIS) | http://www.iobis.org/mapper/?dataset=2 |
| Marine: Body Length | 499 | 0.052 | 95.8 | 15.750 | 58 | 17.5 | 4 | 51.485 | 2.895 | 0.300 | Density | NA | Macrobenthos monitoring at long-term monitoring stations in the Belgian part of the North Sea between 1979 and 1999 | none |
| Marine: Body Length | 246 | 0.063 | 335.0 | 15.000 | 93 | 14.0 | 1 | 25.244 | 121.624 | 0.000 | Count | NA | Long-term monitoring dataset of fish assemblages impinged at nuclear power plants in northern Taiwan | http://datadryad.org/resource/doi:10.506 |
| Marine: Body Length | 252 | 0.067 | 15.8 | 8.308 | 99 | 9.1 | 13 | 32.454 | -78.968 | 0.000 | Count | Weight | MARMAP Blackfish Trap Survey 1990-2009 | http://www2.usgs.gov/obis-usa/search/?datasetid=MARMAP_Blackf |
| Marine: Body Length | 191 | 0.067 | 19.4 | 2.667 | 46 | 8.7 | 56 | 39.123 | -66.641 | 0.000 | Count | NA | NEFSC Benthic Database (OBIS-USA) | http://www.iobis.org/mapper/?dataset=1 |
| Marine: Body Length | 78 | 0.070 | 68.6 | 14.400 | 62 | 19.4 | 10 | 56.730 | 18.236 | 0.000 | Count | NA | IOW Macrozoobenthos monitoring Baltic Sea (1980-2005) (EurOBIS) | http://www.iobis.org/mapper/?dataset=2 |
| Marine: Body Length | 169 | 0.071 | 13.9 | 10.267 | 50 | 16.4 | 791 | 34.858 | -121.615 | 0.900 | Count | NA | CalCOFI and NMFS Seabird and Marine Mammal Observation Data. 1987-2006 (SEAMAP) | http://www.iobis.org/mapper/?dataset=2 |
| Marine: Body Length | 99 | 0.078 | 56.6 | 3.012 | 87 | 9.7 | 256 | -25.598 | 134.025 | 0.000 | NA | Weight | CSIRO Marine Data Warehouse (OBIS Australia) | http://www.iobis.org/mapper/?dataset=5 |
| Marine: Body Length | 232 | 0.081 | 15.1 | 5.615 | 61 | 12.8 | 13 | -56.815 | 93.884 | 0.000 | Count | Weight | Pelagic Fish Observations 1968-1999 | http://gcmd.nasa.gov/KeywordSearch/Me 00038 |
| Marine: Body Length | 85 | 0.090 | 51.5 | 9.906 | 45 | 9.4 | 53 | 53.605 | 4.248 | 0.000 | Density | NA | North Sea observations of Crustacea. Polychaeta. Echinodermata. Mollusca and some other groups between 1986 and 2003 | http://www.emodnet-biology.eu/data-catalog?module=dataset&dasid=1037 |
| Marine: Body Length | 143 | 0.099 | 67.0 | 6.000 | 72 | 7.0 | 1 | -28.494 | -71.860 | 0.000 | Count | NA | COPEPODA-ESPOBIS Data Base IMO-UdeC | http://www.iobis.org/mapper/?dataset=1 |
| Marine: Body Length | 45 | 0.112 | 337.0 | 6.000 | 93 | 5.0 | 1 | -17.500 | -149.000 | 2.000 | Count | Size | MCR LTER Coral Reef Long-term Population and Community Dynamics Fishes | http://mcr.lternet.edu/cgi-bin/showDataset.cgi?docid=knb-lter-mcr.6 |
| Marine: Body Length | 112 | 0.119 | 14.6 | 9.882 | 97 | 14.6 | 907 | 24.981 | -51.374 | 0.000 | Count | NA | NOAA Southeast Fishery Science Center (SEFSC) Commercial Pelagic Observer Program (POP) Data (SEFSC_POP) | http://www.iobis.org/mapper/?dataset=1 |
| Marine: Body Length | 365 | 0.133 | 34.7 | 6.000 | 88 | 5.0 | 3 | -37.256 | 176.325 | 0.025 | Count | NA | Hahei marine dataset (1997-2002) | None |
| Marine: Body Length | 451 | 0.155 | 20.0 | 4.000 | 71 | 7.0 | 1 | 64.000 | -178.000 | 0.000 | NA | Weight | Composition (%) and biomass (thous. tons) of fish species in Gulf of Anadyr in 2005-2012 | http://www.natural-sciences.ru/ru/article/view?id=33118 |
| Marine: Body Length | 211 | 0.178 | 60.2 | 6.750 | 94 | 7.0 | 4 | 32.901 | -78.686 | 0.000 | Count | Weight | MARMAP Fly Net 1990-2009 | http://www.usgs.gov/obis-usa/search/?datasetid=MARMAP_FlyNe |

*(continued)*

| Trait tested | ID | tau | N. Sp. | N. Years | Trait % | Year Span | N. Cells | Latitude | Longitude | Grain | Abundance | Biomass | Study Title | Data Source |
|---|---|---|---|---|---|---|---|---|---|---|---|---|---|---|
| Marine: Body Length | 349 | 0.195 | 78.0 | 3.000 | 49 | 14.0 | 1 | 34.918 | -123.026 | 0.000 | Count | Weight | St. M polychaete species time-series | None |
| Marine: Body Length | 176 | 0.195 | 42.0 | 3.429 | 43 | 7.4 | 107 | 45.073 | -60.564 | 0.000 | Count | Weight | Atlantic Zone Monitoring Program Maritimes Region (AZMP) plankton datasets. In Fisheries and Oceans Canada - BioChem archive (OBIS Canada) | http://www.iobis.org/mapper/?dataset=2 |
| Marine: Body Length | 287 | 0.222 | 13.0 | 5.000 | 58 | 6.0 | 2 | 43.856 | -68.984 | 0.000 | Count | NA | Maine Department of Marine Resources Inshore Trawl Survey 2000?2009 (OBIS-USA) | http://www.iobis.org/mapper/?dataset=1 |
| Marine: Body Length | 297 | 0.226 | 11.3 | 11.000 | 74 | 10.0 | 3 | -17.525 | -149.837 | 0.000 | Count | NA | MCR LTERCoral Reef Long-term Population and Community Dynamics Other Benthic Invertebrates. ongoing since 2005 | http://mcr.lternet.edu/cgi-bin/showDataset.cgi?docid=knb-lter-mcr.7 |
| Marine: Body Length | 171 | 0.226 | 13.3 | 11.167 | 83 | 11.3 | 6 | 24.896 | -76.292 | 0.000 | Count | NA | Bahamas Marine Mammal Research Organisation Opportunistic Sightings (SEAMAP) | http://www.iobis.org/mapper/?dataset=2 |
| Marine: Body Length | 183 | 0.229 | 10.8 | 4.617 | 48 | 9.4 | 68 | 43.777 | -63.751 | 0.000 | Count | NA | DFO Maritimes Research Vessel Trawl Surveys Invertebrate Observations (OBIS Canada) | http://www.iobis.org/mapper/?dataset=2 |
| Marine: Body Length | 450 | 0.242 | 26.0 | 4.000 | 77 | 7.0 | 1 | 60.233 | 168.350 | 0.000 | NA | Weight | Composition (%) and biomass (thous. tons) of fish species in Olyutorsky-Navarin area in 2005-2012 | http://www.natural-sciences.ru/ru/article/view?id=33118 |
| Marine: Body Length | 292 | 0.294 | 23.0 | 2.000 | 93 | 5.0 | 3 | -25.212 | 135.825 | 0.000 | Count | NA | CSIRO Marine Data Warehouse (OBIS Australia) | http://www.iobis.org/mapper/?dataset=5 |
| Marine: Body Length | 452 | 0.309 | 13.0 | 8.000 | 84 | 8.0 | 1 | 47.000 | 160.000 | 0.000 | NA | Weight | Year-to-year dynamics of total nekton biomass (thous. tons) in the upper epipelagic of the North-West Pacific in June-July of 2000s | http://cyberleninka.ru/article/n/vidovaya-struktura-i-mezhgodovaya-dinamika-biomassy-nektona-v-verhney-epipelagiali-prikurilskih-vod-tihogo-okeana-v-letnie-periody-2000-h |
| Marine: Body Length | 67 | 0.333 | 10.0 | 6.000 | 41 | 5.0 | 4 | -28.954 | 24.951 | 0.000 | Count | NA | Animal Demography Unit - Coordinated Waterbird Counts (CWAC) (AfrOBIS) | http://www.iobis.org/mapper/?dataset=6 |
| Plants: Maximum Height | 394 | -0.460 | 10.0 | 3.000 | 100 | 37.0 | 1 | 49.655 | 15.991 | 0.175 | Count | Weight | Zakova hora | http://naturalforests.cz/research |
| Plants: Maximum Height | 398 | -0.457 | 16.0 | 2.000 | 55 | 22.0 | 1 | 49.790 | 15.752 | 0.193 | Count | Weight | Polom | http://naturalforests.cz/research |
| Plants: Maximum Height | 384 | -0.448 | 14.0 | 3.000 | 92 | 33.0 | 1 | 48.655 | 16.941 | 0.173 | Count | Weight | Cahnov-Soutok | http://naturalforests.cz/research |

*(continued)*

| Trait tested | ID | tau | N. Sp. | N. Years | Trait % | Year Span | N. Cells | Latitude | Longitude | Grain | Abundance | Biomass | Study Title | Data Source |
|---|---|---|---|---|---|---|---|---|---|---|---|---|---|---|
| Plants: Maximum Height | 392 | -0.400 | 10.0 | 3.000 | 93 | 39.0 | 1 | 48.880 | 13.836 | 0.162 | Count | Weight | Stozec | http://naturalforests.cz/research |
| Plants: Maximum Height | 487 | -0.368 | 19.0 | 2.000 | 52 | 17.0 | 1 | 78.954 | -77.141 | 0.000 | NA | Cover | ITEX Dataset 9 - Alexfiord (Levdolomite, Levgranite) and Sverdrup | none |
| Plants: Maximum Height | 235 | -0.231 | 18.0 | 4.000 | 72 | 17.0 | 1 | 43.953 | -71.735 | 0.001 | Count | Weight | Forest Inventory of a Northern Hardwood Forest Watershed 5? | http://www.hubbardbrook.org/data/data |
| Plants: Maximum Height | 224 | -0.188 | 50.0 | 3.000 | 67 | 10.0 | 1 | 45.400 | -93.200 | 0.000 | Count | NA | Experiment 133 - Effect of Burning Patterns on Vegetation in the Fish Lake Burn Compartments - Shrub Survey | http://www.cedarcreek.umn.edu/research |
| Plants: Maximum Height | 483 | -0.147 | 54.0 | 24.000 | 49 | 23.0 | 1 | 43.270 | 41.410 | 0.000 | NA | Cover | ITEX Dataset 5 - Teberda (Malaya Alpine-Snowbed and Geranium Hedysarum Meadow) | none |
| Plants: Maximum Height | 492 | -0.130 | 30.0 | 13.000 | 42 | 24.0 | 1 | 68.629 | -149.578 | 0.000 | NA | Cover | ITEX Dataset 14 - Toolik (LTER Heath, LTER Moist acidic tussock, LTER non-acidic tussock, LTER wet sedge, SAG wet sedge2, Tussock 1981 plots) | none |
| Plants: Maximum Height | 401 | -0.085 | 11.0 | 2.000 | 86 | 20.0 | 1 | 49.922 | 13.772 | 0.253 | Count | NA | Kohoutov | http://naturalforests.cz/research |
| Plants: Maximum Height | 214 | -0.068 | 12.4 | 19.583 | 93 | 51.3 | 12 | 45.343 | -122.799 | 0.010 | Count | NA | Long-term growth mortality and regeneration of trees in permanent vegetation plots in the Pacific Northwest 1910 to present | http://andrewsforest.oregonstate.edu/dat |
| Plants: Maximum Height | 486 | -0.055 | 47.0 | 3.000 | 81 | 15.0 | 1 | 46.476 | 9.584 | 0.000 | NA | Cover | ITEX Dataset 8 - Valbercla (Alpine) | none |
| Plants: Maximum Height | 502 | -0.045 | 12.0 | 7.000 | 82 | 47.0 | 1 | 46.920 | -88.026 | 0.001 | Count | Size | Long-term tree demography in old-growth forests of Huron Mts, MI from permanent-plot censuses, 1962-2009 | https://knb.ecoinformatics.org//#view/k |
| Plants: Maximum Height | 389 | -0.042 | 20.0 | 3.000 | 98 | 33.0 | 1 | 48.679 | 16.948 | 0.222 | Count | Weight | Ranspurk | http://naturalforests.cz/research |
| Plants: Maximum Height | 18 | -0.042 | 98.0 | 29.000 | 54 | 50.0 | 1 | 44.330 | -112.330 | 0.000 | Count | NA | Mapped quadrats in sagebrush steppe long-term data for analyzing demographic rates and plant to plant interactions | http://esapubs.org/archive/ecol/E091/24 |
| Plants: Maximum Height | 465 | -0.037 | 31.0 | 6.000 | 84 | 5.0 | 5 | 49.581 | 13.310 | 0.000 | Count | NA | Plants from the Bavarian Forest | None |
| Plants: Maximum Height | 241 | -0.035 | 271.0 | 4.000 | 64 | 13.0 | 1 | 9.364 | -79.955 | 0.060 | Count | NA | Sherman Forest Dynamics Plot Panama | http://www.ctfs.si.edu/site/Sherman/ |

*(continued)*

| Trait tested | ID | tau | N. Sp. | N. Years | Trait % | Year Span | N. Cells | Latitude | Longitude | Grain | Abundance | Biomass | Study Title | Data Source |
|---|---|---|---|---|---|---|---|---|---|---|---|---|---|---|
| Plants: Maximum Height | 396 | -0.020 | 22.0 | 2.000 | 95 | 10.0 | 1 | 49.955 | 14.153 | 0.668 | Count | Weight | Doutnac | http://naturalforests.cz/research |
| Plants: Maximum Height | 495 | -0.019 | 77.0 | 4.000 | 50 | 9.0 | 1 | 62.183 | 1.335 | 0.000 | NA | Cover | ITEX Dataset 17 - Dovre (Kuntshoe) and Faroe (Sornfelli) | none |
| Plants: Maximum Height | 234 | -0.015 | 16.0 | 7.000 | 74 | 37.0 | 1 | 43.953 | -71.740 | 0.001 | Count | Weight | Forest Inventory of a Northern Hardwood Forest Watershed 6 | http://www.hubbardbrook.org/data/datas |
| Plants: Maximum Height | 355 | -0.004 | 324.0 | 33.000 | 60 | 32.0 | 1 | 39.083 | -96.583 | 0.000 | NA | Cover | Plant Species Composition on Selected Watersheds at Konza Prairie | https://portal.lternet.edu/nis/mapbrowse lter-knz.69.9 |
| Plants: Maximum Height | 242 | 0.000 | 12.0 | 7.000 | 72 | 8.0 | 1 | 45.993 | -74.002 | 0.000 | Count | NA | Lac Croche understory vegetation data set (1998 to 2006) | http://esapubs.org/archive/ecol/E088/19 |
| Plants: Maximum Height | 334 | 0.015 | 40.0 | 6.000 | 44 | 18.0 | 1 | 68.630 | -149.576 | 0.000 | NA | Weight | Above ground plant biomass in a mesic acidic tussock tundra experimental site from 1982 to 20000 Toolik lake Alaska | http://arc-lter.ecosystems.mbl.edu/19822000gs81tus |
| Plants: Maximum Height | 307 | 0.017 | 83.0 | 3.000 | 72 | 9.0 | 1 | 42.409 | -85.383 | 0.000 | Count | NA | Kellogg LTER seed bank | http://lter.kbs.msu.edu/datatables/23 |
| Plants: Maximum Height | 60 | 0.075 | 323.0 | 8.000 | 90 | 33.0 | 1 | 9.152 | -79.847 | 0.500 | Count | NA | Forest Census Plot on Barro Colorado Island | http://ctfs.si.edu/ |
| Plants: Maximum Height | 364 | 0.094 | 14.0 | 14.000 | 95 | 17.0 | 1 | 38.901 | -77.069 | 0.000 | NA | Cover | Long-term reductions in anthropogenic nutrients link to improvements in Chesapeake Bay habitat | http://www.pnas.org/content/107/38/165 |
| Plants: Maximum Height | 512 | 0.098 | 71.9 | 2.571 | 87 | 30.0 | 7 | 51.314 | 11.758 | 0.000 | NA | Cover | The Hundt 2001 data - Rudolf Hundt | None |
| Plants: Maximum Height | 480 | 0.106 | 33.0 | 3.000 | 47 | 10.0 | 1 | 61.560 | -135.130 | 0.000 | NA | Cover | ITEX Dataset 2 - Wolfcreek | none |
| Plants: Maximum Height | 496 | 0.118 | 222.0 | 2.000 | 85 | 8.0 | 1 | 46.432 | 8.673 | 0.000 | NA | Cover | ITEX Dataset 18 - Alpine plots (Sonja Wipf) | none |
| Plants: Maximum Height | 489 | 0.143 | 55.0 | 3.000 | 56 | 8.0 | 1 | 60.370 | 7.320 | 0.025 | NA | Cover | ITEX Dataset 11 - Finse (Ridge) | none |
| Plants: Maximum Height | 255 | 0.164 | 22.0 | 10.000 | 76 | 18.0 | 1 | 46.367 | -87.133 | 0.000 | Count | NA | Multi-decade. spatially explicit population studies of canopy dynamics in Michigan old-growth forests | http://www.esapubs.org/archive/ecol/E09 |
| Plants: Maximum Height | 248 | 0.191 | 95.0 | 13.000 | 68 | 12.0 | 1 | 43.833 | -102.167 | 0.000 | Count | NA | Evidence for long-term shift in plant community composition under decadal experimental warming | http://datadryad.org/resource/doi:10.506 |
| Plants: Maximum Height | 481 | 0.195 | 23.0 | 3.000 | 62 | 7.0 | 1 | 73.154 | -79.944 | 0.000 | NA | Cover | ITEX Dataset 3 - Bylot (Mesprairie and Mespolygon) | none |
| Plants: Maximum Height | 298 | 0.203 | 146.0 | 56.000 | 46 | 69.0 | 1 | 38.800 | -99.300 | 0.000 | NA | Cover | Long-term mapped quadrats from Kansas prairie demographic information for herbaceaous plants | http://esapubs.org/archive/ecol/E088/16 |

*(continued)*

| Trait tested | ID | tau | N. Sp. | N. Years | Trait % | Year Span | N. Cells | Latitude | Longitude | Grain | Abundance | Biomass | Study Title | Data Source |
|---|---|---|---|---|---|---|---|---|---|---|---|---|---|---|
| Plants: Maximum Height | 10 | 0.206 | 25.0 | 3.000 | 69 | 12.0 | 1 | 47.400 | -95.120 | 0.000 | Count | NA | Windstorm disturbance without patch dynamics twelve years of change in a Minnesota forest | http://esapubs.org/archive/ecol/E082/01 |
| Plants: Maximum Height | 497 | 0.207 | 59.0 | 29.000 | 47 | 28.0 | 1 | 43.270 | 41.420 | 0.000 | NA | Cover | ITEX Dataset 19 - Teberda (Festuca Varia Grassland, Malaya Alpine Lichen-Heath) | none |
| Plants: Maximum Height | 479 | 0.213 | 75.0 | 9.000 | 56 | 9.0 | 1 | 68.121 | -3.822 | 0.000 | NA | Cover | ITEX Dataset 1 - Abisko (Wet, Dry, Peatland), Kanger (Bashful, Dopey, Sneezy) and Kilpisjarvi | none |
| Plants: Maximum Height | 485 | 0.260 | 128.0 | 3.000 | 85 | 8.0 | 1 | 64.854 | -19.676 | 0.000 | NA | Cover | ITEX Dataset 7 - Akureyri (GA66, MD72, SB63, SY59), Blonduos (SD33, SD34), Dalsmynni (AG4, KD24, KD25), Hjardarland (LH92, SH90), Holtavorduheidi (AH37, AH38, VH49), Modruvellir (LH69, MV51, MV52), Oxnadalsheidi (SA16, SA17, SA19) and Thykkvibaer (HH100, RT81, VE82) | none |
| Plants: Maximum Height | 393 | 0.296 | 15.0 | 3.000 | 100 | 37.0 | 1 | 49.990 | 13.803 | 0.104 | Count | Weight | Velka Ples | http://naturalforests.cz/research |
| Plants: Maximum Height | 243 | 0.314 | 51.0 | 22.000 | 44 | 22.0 | 1 | 37.447 | -75.667 | 0.000 | Count | Cover | Long-term N-fertilized vegetation plots on Hog Island Virginia Coastal Barrier Islands 1992 to 2014 | http://www.vcrlter.virginia.edu/cgi-bin/showDataset.cgi?docid=knb-lter-vcr.106 |
| Fish: Maximum Length | 284 | -0.333 | 14.8 | 2.190 | 52 | 5.2 | 23 | -11.206 | -43.617 | 0.000 | Count | Weight | Previous_fisheries_REVIZEE_Program (Tropical and Subtropical Western South Atlantic OBIS) | http://www.iobis.org/mapper/?dataset=2 |
| Fish: Maximum Length | 428 | -0.223 | 46.0 | 65.737 | 67 | 70.6 | 38 | 58.959 | 9.768 | 0.000 | Count | NA | Long term monitoring of fish abundances from coastal SKagerrak | None |
| Fish: Maximum Length | 119 | -0.204 | 20.5 | 6.017 | 96 | 23.7 | 1896 | 43.987 | -63.670 | 0.000 | Count | Weight | DFO Maritimes Research Vessel Trawl Surveys Fish Observations (OBIS Canada) | http://www.iobis.org/mapper/?dataset=2 |
| Fish: Maximum Length | 271 | -0.140 | 42.5 | 14.500 | 46 | 13.5 | 8 | 34.306 | -119.875 | 0.000 | Count | NA | Santa Barbara Coastal LTER | http://sbc.lternet.edu//index.html |
| Fish: Maximum Length | 191 | -0.106 | 33.5 | 2.000 | 44 | 11.0 | 2 | 39.123 | -66.641 | 0.000 | Count | NA | NEFSC Benthic Database (OBIS-USA) | http://www.iobis.org/mapper/?dataset=1 |
| Fish: Maximum Length | 126 | -0.095 | 52.7 | 4.143 | 47 | 5.3 | 7 | 31.251 | -78.752 | 0.000 | Count | NA | MARMAP Neuston Nets 1990-2009 (OBIS-USA) | http://www.iobis.org/mapper/?dataset=1 |
| Fish: Maximum Length | 507 | -0.083 | 24.9 | 17.500 | 50 | 26.6 | 32 | 31.912 | 34.392 | 445.000 | NA | Weight | Trawl fisheries in Israeli Mediterranean | none |
| Fish: Maximum Length | 180 | -0.081 | 19.4 | 5.252 | 96 | 14.4 | 6328 | 37.771 | -50.793 | 0.333 | Count | NA | ECNASAP - East Coast North America Strategic Assessment (OBIS Canada) | http://www.iobis.org/mapper/?dataset=3 |
| Fish: Maximum Length | 466 | -0.076 | 26.7 | 9.410 | 91 | 15.9 | 156 | 56.991 | -9.073 | 5.000 | Count | NA | Trawl Survey Data from Rockall Scotland (1986 - 2008) | http://www.gov.scot/Topics/marine/scier |

*(continued)*

| Trait tested | ID | tau | N. Sp. | N. Years | Trait % | Year Span | N. Cells | Latitude | Longitude | Grain | Abundance | Biomass | Study Title | Data Source |
|---|---|---|---|---|---|---|---|---|---|---|---|---|---|---|
| Fish: Maximum Length | 123 | -0.071 | 48.6 | 6.885 | 66 | 7.7 | 130 | 43.891 | -68.941 | 0.000 | Count | Weight | Maine Department of Marine Resources Inshore Trawl Survey 2000?2009 (OBIS-USA) | http://www.iobis.org/mapper/?dataset=1 |
| Fish: Maximum Length | 359 | -0.060 | 44.8 | 12.400 | 45 | 11.4 | 6 | 34.309 | -119.874 | 0.000 | Count | NA | SBC LTER Reef Kelp Forest Community Dynamics Fish abundance | https://portal.lternet.edu/nis/mapbrowse lter-sbc.17.27 |
| Fish: Maximum Length | 288 | -0.057 | 13.6 | 5.262 | 92 | 15.3 | 511 | 43.977 | -63.682 | 0.000 | Count | NA | DFO Maritimes Research Vessel Trawl Surveys Fish Observations (OBIS Canada) | http://www.iobis.org/mapper/?dataset=2 |
| Fish: Maximum Length | 505 | -0.057 | 122.1 | 8.133 | 45 | 20.3 | 19 | 32.133 | 34.668 | 445.000 | Count | NA | Fish and marine invertebrates from the Israeli Eastern Mediterranean sea 1990-4, 2000, 2008-2012 | none |
| Fish: Maximum Length | 213 | -0.056 | 39.6 | 9.802 | 83 | 33.4 | 2996 | 36.625 | -72.636 | 0.000 | Count | Weight | Northeast Fisheries Science Center Bottom Trawl Survey Data (OBIS-USA) | http://www.iobis.org/mapper/?dataset=1 |
| Fish: Maximum Length | 190 | -0.051 | 151.3 | 8.667 | 54 | 7.7 | 3 | 17.757 | -64.604 | 0.000 | Count | NA | St. Croix. USVI Fish Assessment and Monitoring Data (2002 - Present) (NOAA-CCMA) | http://www.iobis.org/mapper/?dataset=1 |
| Fish: Maximum Length | 148 | -0.051 | 18.2 | 3.500 | 43 | 5.5 | 132 | -42.407 | 172.260 | 0.000 | NA | Weight | South Western Pacific Regional OBIS Data provider for the NIWA Marine Biodata Information System (South Western Pacific OBIS) | http://www.gbif.org/dataset/83b58bb2-f762-11e1-a439-00145eb45e9a |
| Fish: Maximum Length | 127 | -0.048 | 14.7 | 6.421 | 96 | 6.8 | 19 | 32.335 | -78.992 | 0.000 | Count | Weight | MARMAP Florida Antillean Trap Survey 1990-2009 (OBIS-USA) | http://www.iobis.org/mapper/?dataset=1 |
| Fish: Maximum Length | 212 | -0.047 | 44.6 | 4.162 | 94 | 5.6 | 37 | 31.941 | -78.825 | 0.000 | Count | Weight | MARMAP Yankee Trawl 1990-2009 | http://www.usgs.gov/obis-usa/search/?datasetid=MARMAP_Yanke |
| Fish: Maximum Length | 182 | -0.035 | 16.8 | 14.956 | 51 | 17.4 | 292 | 47.481 | -62.762 | 0.006 | Count | NA | Snow crab research trawl survey database (Southern Gulf of St. Lawrence. Gulf region. Canada) from 1988 to 2010 (OBIS Canada) | http://www.iobis.org/mapper/?dataset=2 |
| Fish: Maximum Length | 467 | -0.035 | 38.6 | 4.000 | 76 | 12.0 | 7 | 40.676 | -8.685 | 0.000 | Count | Weight | Spatial and temporal organisation of a coastal lagoon fish community | None |
| Fish: Maximum Length | 501 | -0.026 | 79.8 | 7.917 | 51 | 6.9 | 12 | 51.530 | 2.730 | 0.000 | Count | NA | Epibenthos and demersal fish monitoring at long-term monitoring stations in the Belgian part of the North Sea | none |
| Fish: Maximum Length | 189 | -0.026 | 175.0 | 9.750 | 52 | 8.8 | 4 | 18.309 | -64.749 | 0.000 | Count | NA | St. John. USVI Fish Assessment and Monitoring Data (2002 - Present) (NOAA-CCMA) | http://www.iobis.org/mapper/?dataset=1 |
| Fish: Maximum Length | 163 | 0.013 | 62.1 | 8.346 | 53 | 9.7 | 1015 | 56.500 | -168.150 | 0.000 | Count | Weight | North Pacific Groundfish Observer (North Pacific Research Board) | http://www.iobis.org/mapper/?dataset=6 |
| Fish: Maximum Length | 135 | 0.033 | 18.0 | 3.500 | 48 | 5.0 | 3 | -13.022 | -45.166 | 0.000 | NA | Weight | Previous_fisheries_REVIZEE_Program (Tropical and Subtropical Western South Atlantic OBIS) | http://www.iobis.org/mapper/?dataset=2 |

*(continued)*

| Trait tested | ID | tau | N. Sp. | N. Years | Trait % | Year Span | N. Cells | Latitude | Longitude | Grain | Abundance | Biomass | Study Title | Data Source |
|---|---|---|---|---|---|---|---|---|---|---|---|---|---|---|
| Fish: Maximum Length | 125 | 0.036 | 19.7 | 7.707 | 93 | 10.1 | 41 | 31.442 | -78.849 | 0.000 | Count | Weight | MARMAP Chevron Trap Survey 1990-2009 (OBIS-USA) | http://www.iobis.org/mapper/?dataset=1 |
| Fish: Maximum Length | 252 | 0.095 | 15.8 | 8.308 | 96 | 9.1 | 13 | 32.454 | -78.968 | 0.000 | Count | Weight | MARMAP Blackfish Trap Survey 1990-2009 | http://www2.usgs.gov/obis-usa/search/?datasetid=MARMAP_Blackf |
| Fish: Maximum Length | 511 | 0.121 | 67.5 | 6.000 | 48 | 10.0 | 2 | -22.821 | -43.149 | 0.000 | Count | Weight | Demersal fish hauls from Guanabara Bay Brazil 2005-2015 | None |
| Fish: Maximum Length | 99 | 0.143 | 14.0 | 2.000 | 57 | 10.0 | 3 | -25.598 | 134.025 | 0.000 | NA | Weight | CSIRO Marine Data Warehouse (OBIS Australia) | http://www.iobis.org/mapper/?dataset=5 |
| Fish: Maximum Length | 211 | 0.182 | 60.2 | 6.750 | 95 | 7.0 | 4 | 32.901 | -78.686 | 0.000 | Count | Weight | MARMAP Fly Net 1990-2009 | http://www.usgs.gov/obis-usa/search/?datasetid=MARMAP_FlyNe |
| Fish: Maximum Length | 112 | 0.210 | 14.6 | 9.881 | 58 | 14.6 | 907 | 24.981 | -51.374 | 0.000 | Count | NA | NOAA Southeast Fishery Science Center (SEFSC) Commercial Pelagic Observer Program (POP) Data (SEFSC_POP) | http://www.iobis.org/mapper/?dataset=1 |
| Fish: Maximum Length | 451 | 0.350 | 20.0 | 4.000 | 81 | 7.0 | 1 | 64.000 | -178.000 | 0.000 | NA | Weight | Composition (%) and biomass (thous. tons) of fish species in Gulf of Anadyr in 2005-2012 | http://www.natural-sciences.ru/ru/article/view?id=33118 |
| Fish: Maximum Length | 450 | 0.371 | 26.0 | 4.000 | 81 | 7.0 | 1 | 60.233 | 168.350 | 0.000 | NA | Weight | Composition (%) and biomass (thous. tons) of fish species in Olyutorsky-Navarin area in 2005-2012 | http://www.natural-sciences.ru/ru/article/view?id=33118 |
| Fish: Maximum Length | 452 | 0.429 | 13.0 | 8.000 | 54 | 8.0 | 1 | 47.000 | 160.000 | 0.000 | NA | Weight | Year-to-year dynamics of total nekton biomass (thous. tons) in the upper epipelagic of the North-West Pacific in June-July of 2000s | http://cyberleninka.ru/article/n/vidovaya-struktura-i-mezhgodovaya-dinamika-biomassy-nektona-v-verhney-epipelagiali-prikurilskih-vod-tihogo-okeana-v-letnie-periody-2000-h |
| Plants: Seed Mass | 394 | -0.511 | 10.0 | 3.000 | 100 | 37.0 | 1 | 49.655 | 15.991 | 0.175 | Count | Weight | Zakova hora | http://naturalforests.cz/research |
| Plants: Seed Mass | 502 | -0.303 | 12.0 | 7.000 | 100 | 47.0 | 1 | 46.920 | -88.026 | 0.001 | Count | Size | Long-term tree demography in old-growth forests of Huron Mts, MI from permanent-plot censuses, 1962-2009 | https://knb.ecoinformatics.org//#view/k |
| Plants: Seed Mass | 486 | -0.262 | 47.0 | 3.000 | 50 | 15.0 | 1 | 46.476 | 9.584 | 0.000 | NA | Cover | ITEX Dataset 8 - Valbercla (Alpine) | none |
| Plants: Seed Mass | 480 | -0.230 | 33.0 | 3.000 | 46 | 10.0 | 1 | 61.560 | -135.130 | 0.000 | NA | Cover | ITEX Dataset 2 - Wolfcreek | none |
| Plants: Seed Mass | 473 | -0.227 | 95.0 | 14.000 | 54 | 13.0 | 1 | 46.317 | -105.800 | 0.000 | Count | NA | Fourteen years of mapped permanent quadrats in a northern mixed prairie | http://esapubs.org/archive/ecol/E092/14 |
| Plants: Seed Mass | 401 | -0.214 | 11.0 | 2.000 | 76 | 20.0 | 1 | 49.922 | 13.772 | 0.253 | Count | NA | Kohoutov | http://naturalforests.cz/research |
| Plants: Seed Mass | 340 | -0.214 | 93.0 | 15.000 | 49 | 14.0 | 1 | 34.296 | -106.927 | 0.000 | Count | NA | Small Mammal Exclosure Study (SMES) Vegetation Data from the Chihuahuan Desert | http://sev.lternet.edu/content/small-mammal-exclosure-study-smes-0 |
| Plants: Seed Mass | 389 | -0.199 | 20.0 | 3.000 | 98 | 33.0 | 1 | 48.679 | 16.948 | 0.222 | Count | Weight | Ranspurk | http://naturalforests.cz/research |

*(continued)*

| Trait tested | ID | tau | N. Sp. | N. Years | Trait % | Year Span | N. Cells | Latitude | Longitude | Grain | Abundance | Biomass | Study Title | Data Source |
|---|---|---|---|---|---|---|---|---|---|---|---|---|---|---|
| Plants: Seed Mass | 234 | -0.162 | 16.0 | 7.000 | 95 | 37.0 | 1 | 43.953 | -71.740 | 0.001 | Count | Weight | Forest Inventory of a Northern Hardwood Forest Watershed 6 | http://www.hubbardbrook.org/data/datas |
| Plants: Seed Mass | 224 | -0.139 | 50.0 | 3.000 | 72 | 10.0 | 1 | 45.400 | -93.200 | 0.000 | Count | NA | Experiment 133 - Effect of Burning Patterns on Vegetation in the Fish Lake Burn Compartments - Shrub Survey | http://www.cedarcreek.umn.edu/research |
| Plants: Seed Mass | 202 | -0.127 | 75.0 | 3.000 | 51 | 12.0 | 1 | 11.599 | 76.534 | 0.500 | Count | NA | Mudumalai Forest Dynamics Plot. India - Smithsonian Tropical Research Institute | http://www.ctfs.si.edu/site/Mudumalai/ |
| Plants: Seed Mass | 235 | -0.067 | 18.0 | 4.000 | 92 | 17.0 | 1 | 43.953 | -71.735 | 0.001 | Count | Weight | Forest Inventory of a Northern Hardwood Forest Watershed 5? | http://www.hubbardbrook.org/data/datas |
| Plants: Seed Mass | 242 | -0.055 | 12.0 | 7.000 | 91 | 8.0 | 1 | 45.993 | -74.002 | 0.000 | Count | NA | Lac Croche understory vegetation data set (1998 to 2006) | http://esapubs.org/archive/ecol/E088/19 |
| Plants: Seed Mass | 298 | -0.040 | 146.0 | 56.000 | 63 | 69.0 | 1 | 38.800 | -99.300 | 0.000 | NA | Cover | Long-term mapped quadrats from Kansas prairie demographic information for herbaceaous plants | http://esapubs.org/archive/ecol/E088/16 |
| Plants: Seed Mass | 512 | -0.034 | 71.9 | 2.571 | 85 | 30.0 | 7 | 51.314 | 11.758 | 0.000 | NA | Cover | The Hundt 2001 data - Rudolf Hundt | None |
| Plants: Seed Mass | 496 | -0.034 | 222.0 | 2.000 | 62 | 8.0 | 1 | 46.432 | 8.673 | 0.000 | NA | Cover | ITEX Dataset 18 - Alpine plots (Sonja Wipf) | none |
| Plants: Seed Mass | 307 | -0.029 | 83.0 | 3.000 | 73 | 9.0 | 1 | 42.409 | -85.383 | 0.000 | Count | NA | Kellogg LTER seed bank | http://lter.kbs.msu.edu/datatables/23 |
| Plants: Seed Mass | 241 | -0.018 | 271.0 | 4.000 | 59 | 13.0 | 1 | 9.364 | -79.955 | 0.060 | Count | NA | Sherman Forest Dynamics Plot Panama | http://www.ctfs.si.edu/site/Sherman/ |
| Plants: Seed Mass | 60 | -0.015 | 323.0 | 8.000 | 70 | 33.0 | 1 | 9.152 | -79.847 | 0.500 | Count | NA | Forest Census Plot on Barro Colorado Island | http://ctfs.si.edu/ |
| Plants: Seed Mass | 214 | -0.008 | 12.4 | 19.583 | 92 | 51.3 | 12 | 45.343 | -122.799 | 0.010 | Count | NA | Long-term growth mortality and regeneration of trees in permanent vegetation plots in the Pacific Northwest 1910 to present | http://andrewsforest.oregonstate.edu/dat |
| Plants: Seed Mass | 495 | -0.004 | 77.0 | 4.000 | 44 | 9.0 | 1 | 62.183 | 1.335 | 0.000 | NA | Cover | ITEX Dataset 17 - Dovre (Kuntshoe) and Faroe (Sornfelli) | none |
| Plants: Seed Mass | 392 | 0.000 | 10.0 | 3.000 | 93 | 39.0 | 1 | 48.880 | 13.836 | 0.162 | Count | Weight | Stozec | http://naturalforests.cz/research |
| Plants: Seed Mass | 398 | 0.022 | 16.0 | 2.000 | 61 | 22.0 | 1 | 49.790 | 15.752 | 0.193 | Count | Weight | Polom | http://naturalforests.cz/research |
| Plants: Seed Mass | 471 | 0.039 | 162.0 | 10.000 | 47 | 9.0 | 1 | 34.350 | -106.880 | 0.000 | Count | NA | Prescribed Burn Effect on Chihuahuan Desert Grasses and Shrubs at the Sevilleta National Wildlife Refuge | http://sev.lternet.edu/data/sev-166 |
| Plants: Seed Mass | 255 | 0.042 | 22.0 | 10.000 | 90 | 18.0 | 1 | 46.367 | -87.133 | 0.000 | Count | NA | Multi-decade. spatially explicit population studies of canopy dynamics in Michigan old-growth forests | http://www.esapubs.org/archive/ecol/E09 |
| Plants: Seed Mass | 364 | 0.048 | 14.0 | 14.000 | 57 | 17.0 | 1 | 38.901 | -77.069 | 0.000 | NA | Cover | Long-term reductions in anthropogenic nutrients link to improvements in Chesapeake Bay habitat | http://www.pnas.org/content/107/38/165 |

*(continued)*

| Trait tested | ID | tau | N. Sp. | N. Years | Trait % | Year Span | N. Cells | Latitude | Longitude | Grain | Abundance | Biomass | Study Title | Data Source |
|---|---|---|---|---|---|---|---|---|---|---|---|---|---|---|
| Plants: Seed Mass | 483 | 0.051 | 54.0 | 24.000 | 47 | 23.0 | 1 | 43.270 | 41.410 | 0.000 | NA | Cover | ITEX Dataset 5 - Teberda (Malaya Alpine-Snowbed and Geranium Hedysarum Meadow) | none |
| Plants: Seed Mass | 355 | 0.053 | 324.0 | 33.000 | 80 | 32.0 | 1 | 39.083 | -96.583 | 0.000 | NA | Cover | Plant Species Composition on Selected Watersheds at Konza Prairie | https://portal.lternet.edu/nis/mapbrowse[illegible] lter-knz.69.9 |
| Plants: Seed Mass | 18 | 0.092 | 98.0 | 29.000 | 61 | 50.0 | 1 | 44.330 | -112.330 | 0.000 | Count | NA | Mapped quadrats in sagebrush steppe long-term data for analyzing demographic rates and plant to plant interactions | http://esapubs.org/archive/ecol/E091/24[illegible] |
| Plants: Seed Mass | 248 | 0.092 | 95.0 | 13.000 | 74 | 12.0 | 1 | 43.833 | -102.167 | 0.000 | Count | NA | Evidence for long-term shift in plant community composition under decadal experimental warming | http://datadryad.org/resource/doi:10.506[illegible] |
| Plants: Seed Mass | 356 | 0.106 | 66.9 | 12.667 | 50 | 36.6 | 9 | -17.038 | 145.561 | 0.005 | Count | NA | Long-term stem inventory data from tropical rain forest plots in Australia | http://esapubs.org/archive/ecol/E095/20[illegible] |
| Plants: Seed Mass | 497 | 0.110 | 59.0 | 29.000 | 45 | 28.0 | 1 | 43.270 | 41.420 | 0.000 | NA | Cover | ITEX Dataset 19 - Teberda (Festuca Varia Grassland, Malaya Alpine Lichen-Heath) | none |
| Plants: Seed Mass | 465 | 0.115 | 31.0 | 6.000 | 79 | 5.0 | 5 | 49.581 | 13.310 | 0.000 | Count | NA | Plants from the Bavarian Forest | None |
| Plants: Seed Mass | 243 | 0.123 | 51.0 | 22.000 | 49 | 22.0 | 1 | 37.447 | -75.667 | 0.000 | Count | Cover | Long-term N-fertilized vegetation plots on Hog Island Virginia Coastal Barrier Islands 1992 to 2014 | http://www.vcrlter.virginia.edu/cgi-bin/showDataset.cgi?docid=knb-lter-vcr.106 |
| Plants: Seed Mass | 485 | 0.142 | 128.0 | 3.000 | 67 | 8.0 | 1 | 64.854 | -19.676 | 0.000 | NA | Cover | ITEX Dataset 7 - Akureyri (GA66, MD72, SB63, SY59), Blonduos (SD33, SD34), Dalsmynni (AG4, KD24, KD25), Hjardarland (LH92, SH90), Holtavorduheidi (AH37, AH38, VH49), Modruvellir (LH69, MV51, MV52), Oxnadalsheidi (SA16, SA17, SA19) and Thykkvibaer (HH100, RT81, VE82) | none |
| Plants: Seed Mass | 336 | 0.157 | 100.0 | 14.000 | 52 | 13.0 | 1 | 31.939 | -109.080 | 0.000 | Count | NA | Long term monitoring and experimental manipulation of a Chihuahuan Desert ecosystem near Portal Arizona | http://esapubs.org/archive/ecol/E090/11[illegible] |
| Plants: Seed Mass | 240 | 0.203 | 167.0 | 13.000 | 56 | 12.0 | 1 | 34.350 | -106.880 | 0.000 | Count | Cover | Pinon-Juniper (Core Site) Quadrat Data for the Net Primary Production Study at the Sevilleta National Wildlife Refuge New Mexico (2003-present ) | http://sev.lternet.edu/node/1718 |

*(continued)*

| Trait tested | ID | tau | N. Sp. | N. Years | Trait % | Year Span | N. Cells | Latitude | Longitude | Grain | Abundance | Biomass | Study Title | Data Source |
|---|---|---|---|---|---|---|---|---|---|---|---|---|---|---|
| Plants: Seed Mass | 489 | 0.208 | 55.0 | 3.000 | 46 | 8.0 | 1 | 60.370 | 7.320 | 0.025 | NA | Cover | ITEX Dataset 11 - Finse (Ridge) | none |
| Plants: Seed Mass | 393 | 0.219 | 15.0 | 3.000 | 100 | 37.0 | 1 | 49.990 | 13.803 | 0.104 | Count | Weight | Velka Ples | http://naturalforests.cz/research |
| Plants: Seed Mass | 384 | 0.231 | 14.0 | 3.000 | 92 | 33.0 | 1 | 48.655 | 16.941 | 0.173 | Count | Weight | Cahnov-Soutok | http://naturalforests.cz/research |
| Plants: Seed Mass | 481 | 0.257 | 23.0 | 3.000 | 47 | 7.0 | 1 | 73.154 | -79.944 | 0.000 | NA | Cover | ITEX Dataset 3 - Bylot (Mesprairie and Mespolygon) | none |
| Plants: Seed Mass | 396 | 0.292 | 22.0 | 2.000 | 91 | 10.0 | 1 | 49.955 | 14.153 | 0.668 | Count | Weight | Doutnac | http://naturalforests.cz/research |
| Plants: Seed Mass | 10 | 0.380 | 25.0 | 3.000 | 79 | 12.0 | 1 | 47.400 | -95.120 | 0.000 | Count | NA | Windstorm disturbance without patch dynamics twelve years of change in a Minnesota forest | http://esapubs.org/archive/ecol/E082/01 |
| Marine: Qualitative Body Size | 367 | -0.135 | 137.0 | 6.000 | 40 | 5.0 | 2 | 34.058 | 134.978 | 0.000 | Count | NA | Monitoring site 1000 Coastal zone research - Tidal flat survey | http://www.biodic.go.jp/moni1000/findin |
| Marine: Qualitative Body Size | 457 | -0.074 | 226.0 | 8.000 | 93 | 11.0 | 1 | 21.945 | 120.769 | 0.000 | MeanCount | NA | Dynamics of a coral reef community | None |
| Marine: Qualitative Body Size | 330 | -0.051 | 80.9 | 4.737 | 57 | 43.1 | 40 | -23.831 | 136.449 | 0.000 | Density | NA | Over 75 years of zooplankton data from Australia | http://esapubs.org/archive/ecol/E095/27 |
| Marine: Qualitative Body Size | 97 | -0.049 | 28.8 | 2.500 | 52 | 17.3 | 6 | 72.739 | 10.694 | 0.000 | Count | NA | Archives of the Arctic Seas Zooplankton (ARC) | http://www.iobis.org/mapper/?dataset=4 |
| Marine: Qualitative Body Size | 191 | 0.044 | 53.5 | 3.529 | 47 | 11.6 | 226 | 39.123 | -66.641 | 0.000 | Count | NA | NEFSC Benthic Database (OBIS-USA) | http://www.iobis.org/mapper/?dataset=1 |
| Marine: Qualitative Body Size | 176 | 0.055 | 45.0 | 4.650 | 52 | 8.0 | 140 | 45.073 | -60.564 | 0.000 | Count | Weight | Atlantic Zone Monitoring Program Maritimes Region (AZMP) plankton datasets. In Fisheries and Oceans Canada - BioChem archive (OBIS Canada) | http://www.iobis.org/mapper/?dataset=2 |
| Marine: Qualitative Body Size | 133 | 0.086 | 28.0 | 2.000 | 81 | 6.0 | 1 | -14.617 | -42.219 | 0.000 | Count | NA | Copepods (Tropical and Subtropical Western South Atlantic OBIS) | http://www.iobis.org/mapper/?dataset=5 |
| Marine: Qualitative Body Size | 143 | 0.133 | 67.0 | 6.000 | 73 | 7.0 | 1 | -28.494 | -71.860 | 0.000 | Count | NA | COPEPODA-ESPOBIS Data Base IMO-UdeC | http://www.iobis.org/mapper/?dataset=1 |
| Marine: Qualitative Body Size | 72 | 0.156 | 35.0 | 2.000 | 46 | 5.0 | 1 | 65.890 | 36.806 | 0.000 | Count | NA | White Sea Plankton | http://www.iobis.org/mapper/?dataset=4 |
